# An allelic series rare variant association test for candidate gene discovery

**DOI:** 10.1101/2022.12.23.521658

**Authors:** Zachary R McCaw, Colm O’Dushlaine, Hari Somineni, Michael Bereket, Christoph Klein, Theofanis Karaletsos, Francesco Paolo Casale, Daphne Koller, Thomas W Soare

**Affiliations:** Insitro, South San Francisco, CA, USA.; Institute of AI for Health, Helmholtz Munich, Neuherberg, Germany. Helmholtz Pioneer Campus, Helmholtz Munich, Neuherberg, Germany. School of Computation, Information and Technology, Technical University of Munich, Munich, Germany.

**Author notes:** Correspondence: ZRM, TWS.

**Keywords:** Allelic series, Rare variant association testing, Variant Pathogenicity, Whole exome sequencing

## Abstract

Allelic series are of candidate therapeutic interest due to the existence of a dose-response relationship between the functionality of a gene and the degree or severity of a phenotype. We define an allelic series as a gene in which increasingly deleterious mutations lead to increasingly large phenotypic effects, and develop a gene-based rare variant association test specifically targeted for the identification of allelic series. Building on the well-known burden and sequence kernel association (SKAT) tests, we specify a variety of association models, covering different genetic architectures, and integrate these into a COding-variant Allelic Series Test (COAST). Through extensive simulations, we confirm that COAST maintains the type I error and improves power when the pattern of coding-variant effect sizes increases monotonically with mutational severity. We applied COAST to identify allelic series for 4 circulating lipid traits and 5 cell count traits among 145,735 subjects with available whole exome sequencing data from the UK Biobank. Compared with optimal SKAT (SKAT-O), COAST identified 29% more Bonferroni significant associations with circulating lipid traits, on average, and 82% more with cell count traits. All of the gene-trait associations identified by COAST have corroborating evidence either from rare-variant associations in the full cohort (Genebass, *N* = 400K), or from common variant associations in the GWAS catalog. In addition to detecting many gene-trait associations present in Genebass using only a fraction (36.9%) of the sample, COAST detects associations, such as *ANGPTL4* with triglycerides, that are absent from Genebass but which have clear common variant support.

## Introduction

The term *allelic series* refers to a collection of alleles in a gene or pathway that leads to a gradation of possible phenotypes [1]. Genes demonstrating allelic series are of candidate therapeutic interest due to the existence of a dose-response relationship between the functionality of the gene and the magnitude or severity of the phenotype [2]. This relationship provides evidence in humans that pharmacological modulation of the implicated gene has the potential to move the target phenotype in the direction of clinical benefit. Seminal work by Plenge *et al* [2] highlighted the opportunity for identifying such dose-response relationships preclinically from the natural genetic variation found in human populations. Such ‘experiments of nature’ were instrumental to the development of statins [2], and have confirmed the role of *PCSK9* in familial hypercholesterolaemia, LDL cholesterol, and coronary artery disease, as reviewed in [3]. Another impactful example is the allelic series in *TYK2* [4], now the basis for a successful selective inhibitor (deucravacitinib) for the treatment of psoriasis [5]. From the standpoint of drug design, the advantage of focusing on allelic series is that modulation of the gene has already been linked with variation in phenotype, reducing the risk that an intervention targeting that gene will fail for lack of efficacy. Motivated by this paradigm, we introduce a rare variant association test intended for identifying allelic series in large genotype-phenotype cohorts, such as the UK Biobank [6]. Hereafter, by allelic series, we refer specifically to an individual gene wherein increasingly deleterious mutations have increasingly large phenotypic effects.

Genome-wide association studies (GWAS) canonically seek to associate common genetic variants, typically those having a minor allele frequency (MAF) exceeding 1 – 5%, with complex traits and diseases [7]. The advent of whole genome and whole exome sequencing studies has enabled rare variant association analysis, wherein associations are sought for variants having lower MAFs [8, 9]. Because even large cohorts may have few examples of such subjects, specialized methodology is required to reliably associate phenotypes with rare variants [10]. For a given effect size, the power of single-variant association tests declines precipitously with the MAF [11]. To overcome this limitation, rare variant association tests typically aggregate signal within a biologically meaningful region, such as a gene.

Two major subdivisions of rare variant association tests are burden tests and variance component tests. Burden tests [12, 13, 14] associate the phenotype with a single genetic score formed by pooling the variants in a region. These tests can differ with respect to the assignment of weights to variants but share in common the assumption that all rare variants in the region affect the phenotype in the same direction [10]. When this is not the case, the power of burden testing is diminished. Variance component tests, notably the sequence kernel association test (SKAT) [15], relax the assumption of a common direction of effect by instead supposing the effect sizes are random, following a distribution with mean zero and finite variance. SKAT performs well when some of the rare variants in the region are non-causal or have conflicting directions of effect, but loses power when the genetic architecture is consonant with that assumed by the burden test. In practice the genetic architecture of a trait is seldom known. Optimal SKAT (SKAT-O) [16] adaptively combines the SKAT and burden tests to provide high power for rare variant association testing under the scenarios assumed by either the component tests.

There are several existing approaches that incorporate functional annotations to generally improve power for gene-based association testing [17, 18, 19]. For example, STAAR [19] uses annotation principal components to upweight the probability of a variant being causal based on the fractional rank of that variant’s annotation score, relative to all sequenced variants, within functional categories such as epigenetic marks, conservation, and protein function. In a different vein, Regenie [20] and STAARpipeline [21] allow for the definition of masks that specify which variants to include in an association test based on functional categories. However, no existing methods are specifically targeted to the identification of allelic series. We fill this gap by defining a rare variant association test that incorporates a set of adjustable allelic series weights to encourage rejection of the null hypothesis in cases where the expected effect of a variant on the phenotype increases monotonically with the severity of the mutation. Our approach focuses on the rare coding variants within a gene, as annotated by the Ensembl Variant Effect Predictor (VEP) [22]. Specifically, our test operates on three classes of variants: the benign missense variants (BMVs), deleterious missense variants (DMVs), and protein truncating variants (PTVs). Building on the burden and SKAT tests, we define a variety of gene-based association models targeting different genetic architectures. These tests are aggregated into a single, omnibus, COding-variant Allelic Series Test (COAST). In this work, we present the first rare variant association test specifically intended for the identification of genes harboring a dose-response relationship with the phenotype, and highlight its potential for drug discovery by demonstrating its efficacy for uncovering allelic series. Through extensive simulation studies, we validate that COAST controls the type I error in the absence of a genotype-phenotype association, and benchmark its power relative to SKAT-O [10] under a variety of genetic architectures. We select SKAT-O as the comparator due to its widespread use, computational efficiency, and direct applicability using only the data required by COAST. We note, however, that the set of genes targeted by COAST is narrower than that targeted by SKAT-O, and we are aware of no existing association test that specifically targets allelic series. Using whole exome sequencing data on up to 145,735 subjects and up to 17,225 genes from the UK Biobank, we apply COAST to identify allelic series for 4 circulating lipid traits and 5 cell count traits. Compared with SKAT-O, COAST identified 29% more Bonferroni significant associations with circulating lipids, on average, and 82% more with cell counts. Notably, all of the additional genes detected by COAST have supporting evidence from published common or rare variant association studies, however not all of the rare variant associations have previously been identified by SKAT-O applied to the full UK Biobank (Genebass) [23].

## Material and Methods

### Setting

Let *Y* denote a quantitative phenotype (e.g. circulating cholesterol or erythrocyte count), *X* a set of covariates (e.g. age, sex, genetic principal components), and *G* = (*G*_1_, · · ·, *G_J_*) ∈ {0, 1, 2}*^J^* the additively coded genotype at the *J* rare coding variants within a given gene. To each variant *j*, assign a categorical functional annotation *A_j_* ∈ {1, · · ·, *L}*. For the present work, we specialize to the following *L* = 3 VEP categories:

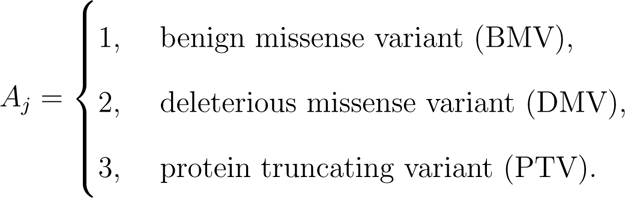

For a given subject, let *N_l_* count the total number of category *l* alleles present in the gene:

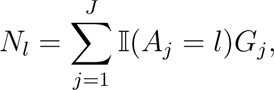

and let *N* = (*N*_1_, *N*_2_, *N*_3_) denote the vector of category-level allele counts.

### Models

#### Standard Burden and SKAT Tests

Consider the standard gene-level association model:

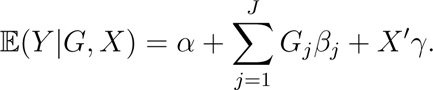

The marginal score for evaluating *H*_0_ : *β_j_* = 0 is 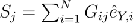 where *i* indexes subjects and 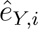 is the residual from regressing *Y* on *X* (but not *G*). Following [10], the standard burden and SKAT tests are expressible as:

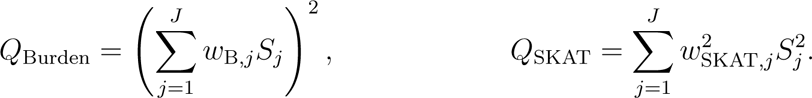

Different choices for the per-variant burden *w*_B_*_,j_* and SKAT *w*_SKAT_*_,j_* weights provide differing powers to detect an association. The standard burden test assigns a weight of *w*_B_*_,j_* = 1 to all variants within a class of interest (e.g. to PTVs), whereas the standard SKAT test [15] assigns weights from the beta distribution based on the variant’s minor allele frequency (MAF): 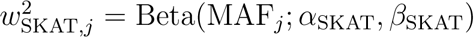. Selecting *α*_SKAT_ = *β*_SKAT_ = 1*/*2 gives rise to inverse variance weighting 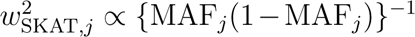, reflecting a genetic architecture in which rare variants have larger effects. Finally, optimal SKAT (SKAT-O) is based on a convex combination of the burden and SKAT statistics [16]:

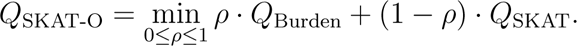

Here *ρ* ∈ [0, 1] adaptively combines *Q*_Burden_ and *Q*_SKAT_.

#### Baseline Model

The **baseline allelic series model** takes the form:

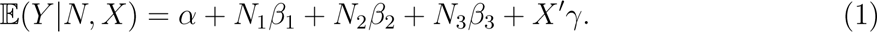

(1) is reminiscent of the standard burden model but differs in that aggregation is grouped by annotation category and a separate coefficient is allocated to each. As such, the baseline allelic series model is not a case of the standard burden test. (1) is also a count-based model, where the expected value of the phenotype increases linearly by *β_l_* for each additional category *l* allele. An indicator-based model replaces the category *l* allele count *N_l_* by an indicator that *N_l_*is non-zero:

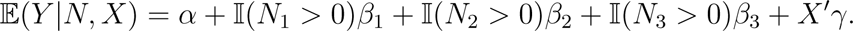

The indicator-based model posits a genetic architecture in which the presence versus absence of category *l* alleles has an effect on the phenotype, but beyond the first, the number of category *l* alleles is immaterial. For example, the presence of any PTVs in a gene may abrogate its function, but additional PTVs beyond the first may have no effect. Under both the count and indicator models, the null hypothesis that the rare coding variants within the gene are unrelated to the phenotype is evaluated by testing *H*_0_ : *β*_1_ = *β*_2_ = *β*_3_ = 0 using a standard Wald or score test with 3 degrees of freedom [24]. Details of the Wald and score tests are provided in the Supplemental Methods.

#### Allelic Series Sum Model

The baseline model provides a test for identifying genes associated with the phenotype and allows variants belonging to different annotation categories to have differing effects. However, it is not yet tailored to the identification of allelic series. We next introduce a set of adjustable **allelic series weights** *w* = (*w*_1_, …, *w_L_*), one for each variant category, in order to specify the *pattern* of effect sizes sought. Any monotone increasing pattern corresponds to an allelic series, but as a simple default, we adopt *w* = (1, 2, 3). We define the **allelic series sum model** as a specialization of (1) under the constraint that *β_l_* = *w_l_β*:

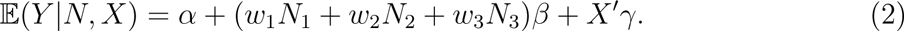

(2) continues to allow different categories to have differing effects, but the constraint requires that the effect size vector (*β*_1_, *β*_2_, *β*_3_) is proportional to the allelic series weights:

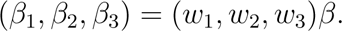

Compared with (1), (2) reduces the number of free parameters from 3 to 1. The null hypothesis of no association between the gene and the phenotype is evaluated by testing *H*_0_ : *β* = 0. The allelic series sum model is a case of the standard burden model with customized weights: 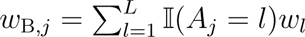. The baseline model may outperform the sum model for architectures where the pattern of effect size differs markedly from that sought by *w*, e.g. (*β*_1_, *β*_2_, *β*_3_) ∞ (0, 0, 1). Conversely, the sum model will have better power the more nearly the prespecified allelic series weights (*w*_1_, *w*_2_, *w*_3_) are proportional to the true pattern of effect sizes (*β*_1_, *β*_2_, *β*_3_).

#### Allelic Series Max Model

Focusing on the parenthetical term in (2), we see that the allelic series sum model aggregates the per-category allele counts (*N*_1_, *N*_2_, *N*_3_) by taking a weighted sum. By using different methods of aggregation, we can construct additional allelic series tests. The **allelic series max model** aggregates via the weighted maximum:

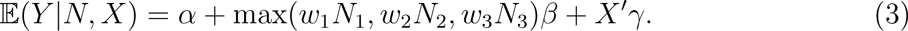

(3), particularly the indicator version, is motivated by a genetic architecture in which the expected phenotype is determined by the most deleterious variant present. For example, the presence of a BMV may have negligible phenotypic impact in the presence of a PTV. As for the sum model, the null hypothesis of no association is evaluated by testing *H*_0_ : *β* = 0. The allelic series max model is not readily expressible within the standard burden framework.

#### Allelic Series SKAT Model

The preceding tests make the burden-type assumption that all variants affect the phenotype in the same direction. The sequence kernel association test (SKAT) [15] was introduced to allow variants to have differing directions of effect. Briefly, SKAT posits that each variant *G_j_* has a separate effect *β_j_* and that these effects are drawn at random from an unspecified distribution with mean zero and variance 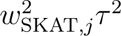, where *τ*^2^ is a common variance component. The null hypothesis of no association is evaluated by testing *H*_0_ : *τ*^2^ = 0. To develop an allelic series test that allows rare variants to have differing directions of effect, we build on the existing SKAT framework but modify the SKAT weights to incorporate the allelic series weights:

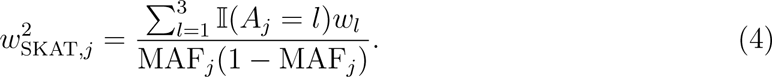

Under (4), the variance of the distribution from which *β_j_* is drawn, and thus the expected magnitude of effect, is directly proportional to *w_l_* and inversely proportional to MAF*_j_*. In the allelic series setting where *w_l_* increases monotonically from BMVs to DMVs to PTVs, this encodes the expectation that rare and more deleterious variants will have larger effects on the phenotype.

#### Allelic Series Omnibus Test

We have now defined several burden-type tests and a SKAT-based test. Which test is best-powered for a particular gene-phenotype pair will depend upon which best approximates the true genetic architecture. Moreover, for a given phenotype, a single test is unlikely to be optimal for all genes, as the genetic architecture can vary from gene to gene. To obviate the need for deciding among tests, we apply each of the preceding then combine their p-values via the Cauchy combination method introduced in [25], a strategy which has also been applied by ACAT-V [26], STAAR [19], and SAIGE-GENE+ [27]. Given *M* p-values (*p*_1_, …, *p_M_*), the **omnibus test statistic** is defined by:

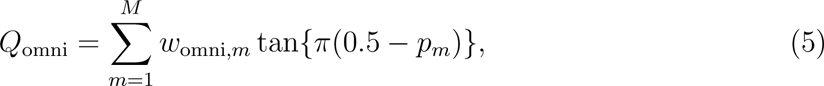

where the omnibus weights *w*_omni_*_,m_* are selected to give burden-type and SKAT-type tests equal weight. Finally, the omnibus p-value is given by:

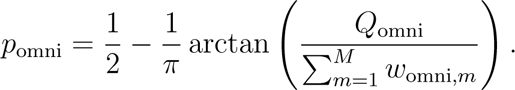

We refer to the omnibus test and its associated p-value as the COding-variant Allelic Series Test (COAST). By default, COAST includes *M* = 7 tests: 6 allelic series burden tests ({base, sum, max} × {count, indicator}) with weight *w*_omni_*_,m_* = 1*/*12 and 1 allelic series SKAT test with weight *w*_omni_*_,m_* = 1*/*2. Optionally, SKAT-O applied to all variants or SKAT-O applied to PTVs only may be included in the omnibus test. Using an omnibus statistic makes COAST robust in the sense that it will have power to detect an association if any of the component models is powered to detect the association [26].

### Simulation Methods

Our simulation studies consider sample sizes of 1K, 5K, 10K, and 50K. Rare variants in linkage equilibrium were simulated to have a minimum minor allele count of 1 and a maximum empirical minor allele frequency of 1%. Variants were assigned annotations of BMV, DMV, or PTV in a 5:4:1 ratio to emulate the empirical frequencies of BMVs, DMVs, and PTVs across all 17,225 genes in our UK Biobank analysis (see below), which were 51.6%, 39.9%, and 8.5% respectively. Simulation studies were also conducted using real *APOB* genotypes and VEP annotations from randomly selected UK Biobank participants. For each subject, covariates were simulated representing age, sex, and 3 genetic principal components (PCs); 10% of phenotypic variation was explained by age, 10% by sex, and 20% by the 3 genetic PCs collectively. For type I error simulations, genotype had no effect on the phenotype. For power simulations, phenotypes were generated under multiple genetic architectures. Each simulated gene contained 10^2^ variants. A random subset of these variants, between 25% and 100%, were selected as causal. The effect size of non-causal variants were zeroed out. Burden phenotypes were simulated from the {baseline, sum, max} models, varying the generative effect sizes (*β*_1_, *β*_2_, *β*_3_). The sum and max phenotypes were simulated by fixing *β* = 1 and varying the generative weights (*w*_1_, *w*_2_, *w*_3_). For SKAT phenotypes, the effect size of variant *j* was generated as: 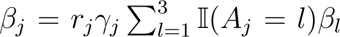. Here *r_j_* is a random sign, 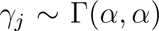 is a positive random scalar with 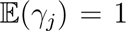 and 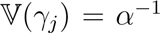 (*α* = 1 for simplicity), and the term 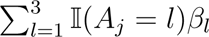 incorporates an additional fixed scalar that depends on the variant’s annotation. Note that the sum and max architectures are not compatible with a SKAT phenotype.

### Preparation of UK Biobank Data

#### Whole Exome Sequencing Data

Variants in the UKBB WES variant callset (N=200K) were filtered out if they were located > 100 bp away from the exome capture regions, located within the ENCODE black list [28], or within low-complexity regions [29]. We further filtered to autosomal, biallelic variants meeting the following criteria: allele balance fractions of ≤ 0.1 (homozygous) or ≥ 0.25 (heterozygous), call rate ≥ 0.95, mean depth (DP) 12 ≤ DP < 200, mean genotype quality GQ ≥ 20, Hardy-Weinberg equilibrium *p >* 1 × 10^−6^, and inbreeding coefficient *F > −*0.3. Variants were annotated with VEP v96 [22] using the following transcript consequences (for any transcript of a gene): protein-truncating variants (PTV): splice acceptor variant, splice donor variant, stop gained, or frameshift variant; missense: missense variant, inframe deletion, inframe insertion, stop lost, start lost, or protein altering variant. Missense variants were annotated to be damaging (DMV) if predicted to be probably damaging or possibly damaging by PolyPhen2 [30] and deleterious or deleterious low confidence by SIFT [31]. Missense variants were annotated to be benign (BMV) if predicted to be benign by PolyPhen2 and tolerated or tolerated low confidence by SIFT [32, 33]. PTVs were further filtered to remove low confidence loss-of-function by LOFTEE [34].

#### Quality Control

Sample-level quality control (QC) was performed using 104,616 high-confidence variants passing the above filtering criteria, and had minor allele count of ≥ 5, and were LD-pruned to *r*^2^ < 0.05. To mitigate confounding due to population structure, samples were first filtered to unrelated individuals without sex chromosome aneuploidy, within ±7 standard deviations of the mean of the first six genotype principal components [35], self-reported ethnic background of White British, Irish, or White, had call rate ≥ 0.95, mean DP ≥ 19, and mean GQ ≥ 47.5. We regressed the top 20 genetic principal components from the following sample QC metrics and removed samples that were > 4 median absolute deviations (MAD) from the median for any of the following metrics: number of SNPs, number of heterozygous variants, number of homozygous alternate variants, number of insertions, number of deletions, number of transitions, number of transversions, the ratio between the number of transitions and transversions, the ratio between the number of heterozygous and homozygous alternate variants, and the ratio between the number of insertions and deletions [34]. The sample size after filtering was 145,753. All data preprocessing was performed in hail 0.2.37 [36].

#### Association Analysis

Processed WES data were restricted to BMVs, DMVs, and PTVs with a sample minor allele frequency ≤ 1%. Following [25, 27], ultra-rare variants, those with a sample MAC ≤ 10, were collapsed into a single pseudo marker separately for each gene × variant category. Genes were required to have at least 3 distinct rare variants (including the pseudo marker) for inclusion in the genome-wide screen; 17,225 genes passed this threshold. The circulating lipid phenotypes analyzed included low and high density lipoprotein (LDL and HDL; UKB Field IDs 30780 and 30760), triglycerides (30870), and total cholesterol (30690). The cell count phenotypes included erythrocytes (30010), leukoctyes (30000), lymphocytes (30120), neutrophils (30140), and thrombocytes (30080). In all cases, the phenotype was transformed to normality by applying the rank-based inverse normal transformation (INT) [37]. Covariates included age (to degree 3), genetic sex, age × sex interaction (to degree 3), 20 genetic principal components, and an indicator for being among the first 50K exomes sequenced. Association analyses were performed using the COAST function from the AllelicSeries R package (v0.0.2) and the SKAT function from the SKAT R package (v2.2.5).

### Overlap Analysis

For each trait of interest, common-variant associations were obtained from the NHGRI-EBI GWAS Portal [38, 39] and filtered to those having *p <* 5 × 10^−8^. Rare-variant associations from both putative loss of function (pLoF) variants and missense variants, including low-confidence pLoF variants and in-frame insertions or deletions, were obtained from the Genebass Portal [23, 40]. Associations were filtered to those Bonferroni significant based on the number of genes with non-missing p-values, and the union of significant associations from the pLoF and missense analyses was taken.

## Results

### Simulation Studies

Simulation studies were performed to evaluate the type I error (validity) and power of the proposed COding-variant Allelic Series Test (COAST). Across sample sizes ranges from 1K to 50K, the p-values from COAST were uniformly distributed under the null hypothesis of no association (Panel A of **Figure 1**). The p-values were uniformly distributed both as applied to real genotypes from the *APOB* gene (133 variants: 79 BMVs, 48 DMVs, 6 PTVs; **Figure 1, Supplemental Figure 1**) and as applied to simulated genotypes in linkage equilibrium (BMVs, DMVs, PTVs in a 5:4:1 ratio; **Supplemental Figure 2**). Empirical demonstration that the type I error is controlled genome-wide is provided by the analyses of permuted phenotypes in **Supplemental Figures 8 and 12**. Across sample sizes, the distribution of p-values from COAST was comparable to that of SKAT-O (**Supplemental Figure 3**). Expected *χ*^2^ statistics for COAST and its components are presented in **Supplemental Figure 4**. COAST is slightly conservative for larger p-values (which are not of interest) but well calibrated in the tails, a known consequence of Cauchy combination being more accurate for smaller p-values [25]. Overall, these findings support the validity of COAST, and its components, for genome-wide analyses.

**Figure 1:**
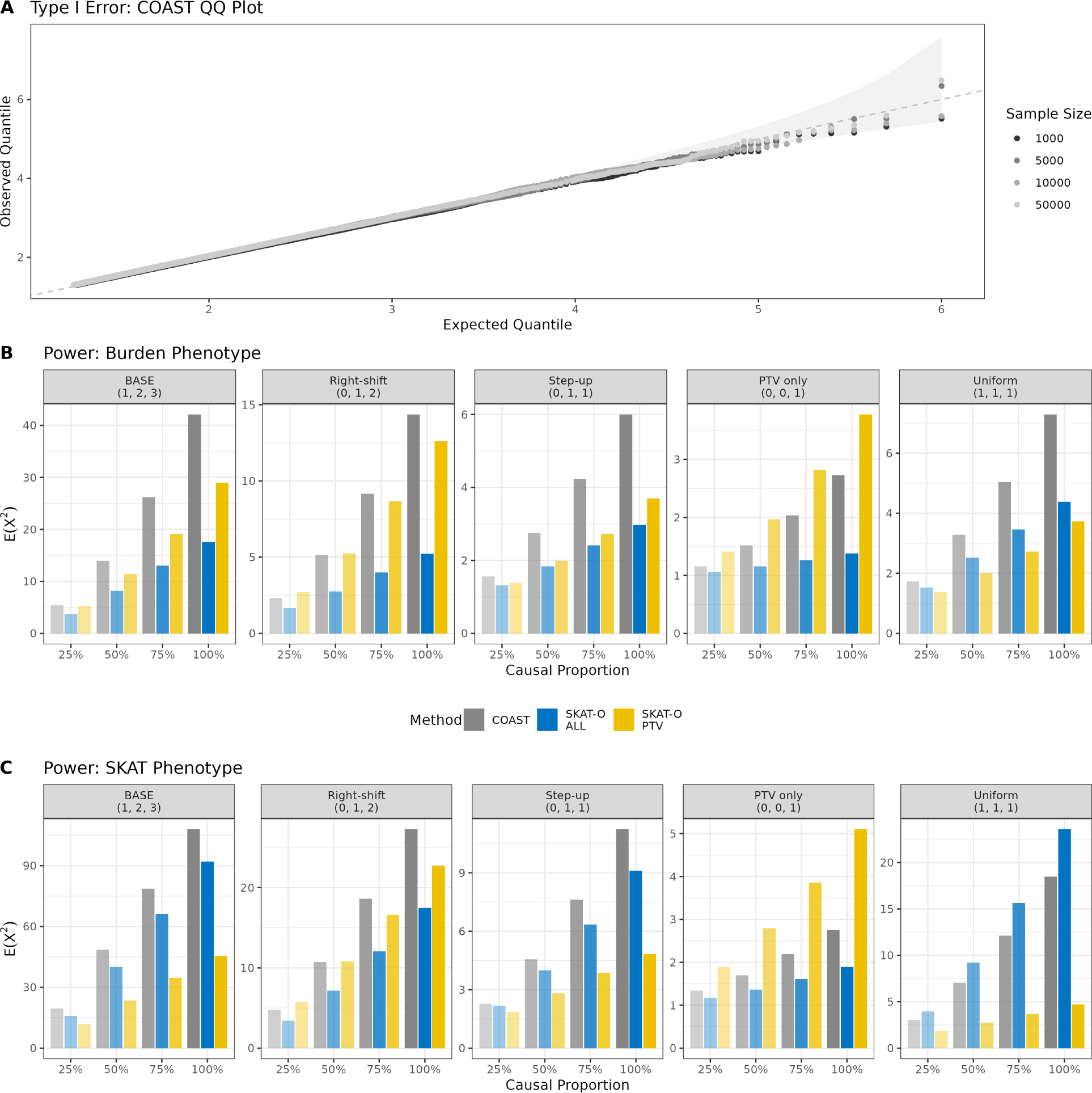
Type I Error and Power of the Coding Allelic Series Test. **A:** Observed versus expected quantiles of − log_10_(*p*) for COAST applied to null phenotypes at various sample sizes, using real genotypes from the *APOB* gene. Adherence to the dashed 45*^◦^* line indicates that the observed p-values are uniformly distributed. Results from 10^6^ simulation replicates at each sample size. **B & C:** Power across various genetic architectures at sample size *N* = 10^4^ using simulated genotypes. Genes included 10^2^ variants with a BMV:DMV:PTV ratio of 5:4:1. Each subplot corresponds to a different genetic architecture generated via (1). The tuple in the heading denotes the relative effect sizes (*β*_1_, *β*_2_, *β*_3_) of BMVs, DMVs, and PTVs. The allelic series weights remained fixed at *w* = (1, 2, 3). For the burden phenotype, the effect sizes of all variants were in the same direction, whereas for the SKAT phenotype, the magnitude and direction of effect were randomized. Each bar aggregates results from 10^4^ simulation replicates.

Panels B and C of **Figure 1** compare the power to detect allelic series between COAST and SKAT-O applied either to all variants (SKAT-O ALL; including BMVs, DMVs, PTVs) or applied to PTVs only (SKAT-O PTV). The weights of COAST were fixed at *w* = (1, 2, 3). Various genetic architectures are generated from (1) with different choices for (*β*_1_, *β*_2_, *β*_3_), as indicated by the tuple in each panel heading. Note that, for all architectures save BASE, COAST is performing with misspecified weights.

For all methods, power to detect an association increased with the proportion of causal variants in the gene. However, the rank ordering of tests by power was insensitive to the causal proportion. For the burden phenotype in Panel B, COAST was uniformly more powerful than SKAT-O ALL, and only superseded by SKAT-O PTV in the case of a PTV-only architecture. Intuitively, the power advantage of COAST declined with the *L*_1_ distance of the generative effect sizes from those specified by the allelic series weights (**Supplemental Figure 7**). For the SKAT phenotype in Panel C, COAST was most powerful in cases of allelic series, but SKAT-O PTV performed best in the case of a PTV-only phenotype, and SKAT-O ALL performed best in the case of a uniform phenotype, meaning the variant annotation was uninformative as to the expected effect size. It is noteworthy that the only architectures for which COAST does not achieve the best power are those that do not exhibit a dose-response relationship (as indicated by the weight tuple). Overall, these findings suggest that although COAST loses some power when its weights do not match the true pattern of effective sizes, COAST is generally well-powered for the task of detecting allelic series.

### Application to Circulating Lipid Phenotypes

As our first case study, we applied COAST and SKAT-O to identify genes associated with the following circulating lipid phenotypes: cholesterol, HDL, LDL, and triglycerides. **Figure 2** presents the number of Bonferroni significant genes identified by COAST and SKAT-O. On average, COAST identified 29% more significant associations than SKAT-O ALL, and 3.3-times as many associations as SKAT-O PTV (**Supplemental Table 1**). Across the union of genes declared significant by any association test, the average *χ*^2^ statistic of COAST was 24% higher than that of SKAT-O ALL and 2.3-times higher than that of SKAT-O PTV (**Supplemental Table 2**). Among 43 total associations, 9 (21%) were unique to COAST, 33 (77%) were in common, and 1 (2%) was unique to SKAT-O ALL (**Supplemental Figure 9**). The uniform quantile-quantile plots in **Supplemental Figure 8** evince no inflation of the type I error.

**Figure 2:**
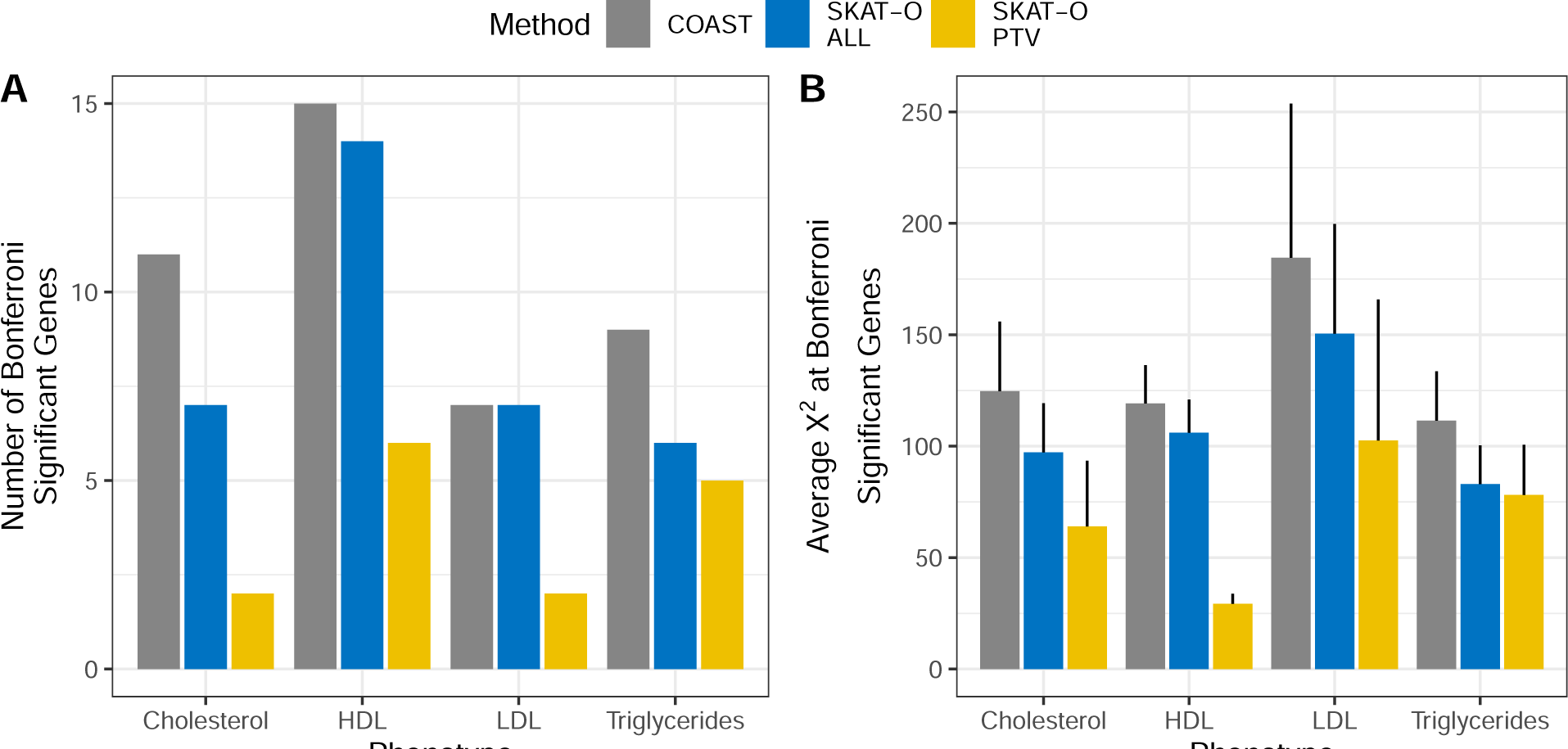
Number and significance of genes significantly associated with circulating lipid phenotypes. **A.** Number of significantly associated genes by phenotype and association test. **B.** Average *χ*^2^ across the union of genes considered Bonferroni significant by any association test. Error bars are 95% confidence intervals.

Taking cholesterol as an example, a down-sampling analysis was performed in which nested subsets of 25%, 50%, and 75% of the total sample were selected, and association analysis was performed on each (**Supplemental Figure 10**). At all sample sizes, COAST identified more Bonferroni significant associations than SKAT-O. Using only 50% of the sample, COAST recovered the same number of associations as SKAT-O ALL in the full sample.

The mirrored Manhattan plots in **Figure 3** show that the signals identified by COAST and SKAT-O all are generally concordant, with several additional associations crossing the Bonferroni threshold under COAST. Manhattan plots for COAST stratified by trait are available in **Supplemental Figure 11**. **Supplemental Tables 3-4** assess the extent to which the gene-trait associations identified by COAST and SKAT-O have existing support from rare variants (Genebass) or common variants (GWAS Catalog). All gene-trait associations had support from at least one of Genebass or the GWAS Catalog, and most had support from both. Many well-known gene-trait associations appear as positive controls, for example *APOB* (*p* = 3.8 × 10^−185^) and *PCSK9* (*p* = 5.5 × 10^−68^) with LDL, *ABCA1* (*p* = 1.3 × 10^−121^) with HDL, and *ANGPTL3* (1.0 × 10^−54^) with triglycerides. It should be noted that while most of the associations reported here are already present in Genebass, the sample size of this analysis is only 36.9% that of Genebass (145, 735 vs. 394, 841). The two genes, both associated with triglycerides, that were identified by COAST and were not present in Genebass were *ANGPTL4* (*p* = 6.7 × 10^−14^) and *A1CF* (*p* = 3.4 × 10^−7^).

**Figure 3:**
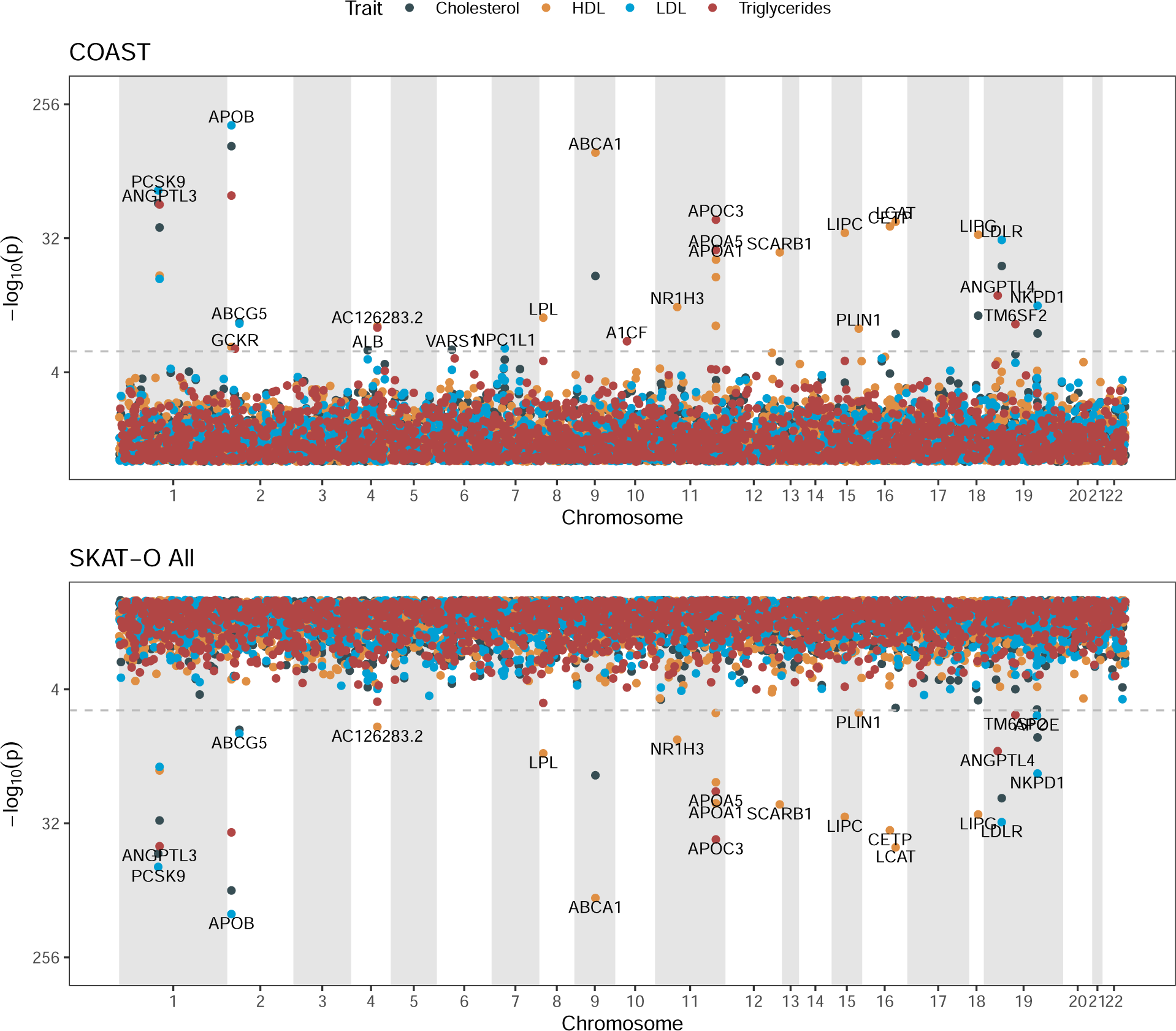
Mirrored Manhattan plots for lipid phenotypes. Upper panel is the coding allelic series test (COAST). Lower panel is SKAT-O applied to all rare coding variants (SKAT-O ALL).

**Figure 4** presents the pattern of effect sizes among genes significantly associated with lipid phenotypes by either COAST or SKAT-O. With few exceptions, the mean effect size magnitude increases monotonically from BMVs to DMVs to PTVs (e.g. *PCSK9* with cholesterol). The two genes associated with triglycerides by COAST that were not present in Genebass (*ANGPTL4* and *A1CF*) both had a monotone increasing pattern of effect sizes. The one gene (*APOE*) associated with LDL by SKAT-O but not COAST had an anti-allelic series pattern, meaning that the effect size magnitude decreased monotonically from BMVs to DMVs to PTVs. COAST retains some power to identify genes that constitute a partial allelic series. For example, the association of *LDLR* with cholesterol was significant, the pattern of effect sizes for BMVs and DMVs increasing as expected, but the effect size for PTVs deviating from expectations. We return to this point in the discussion.

**Figure 4:**
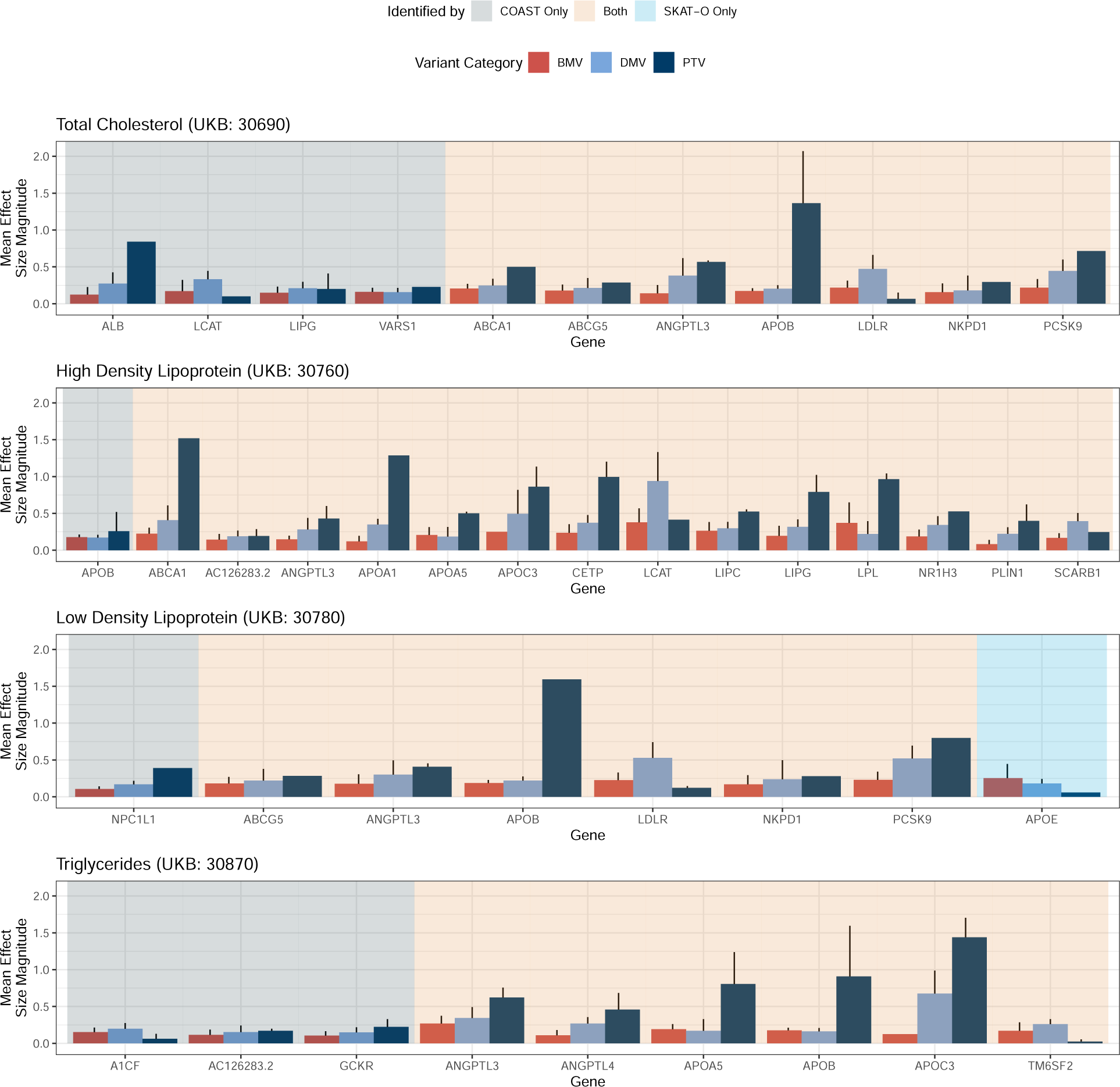
Effect size patterns among genes significantly associated with lipid traits. Effect sizes were estimated by standard linear regression. Each bar represents the mean effect size for variants within a gene and variant category. Error bars are 95% confidence intervals. The absence of an error bar indicates insufficiently many variants to estimate a standard error.

### Application to Cell Count Phenotypes

As a second case study, we applied COAST and SKAT-O to a collection of cell count phenotypes: erythrocytes, leukoctyes, lymphocytes, neutrophils, and thrombocytes (platelets). The uniform quantile-quantile plots in **Supplemental Figure 12** again demonstrate control of the type I error. For all cell count traits, COAST identified more Bonferroni significant associations than SKAT-O (**Figure 5**): 82% more than SKAT-O ALL and 6.3-times as many as SKAT-O PTV (**Supplemental Table 5**). Across the union of genes declared significant by any association test, the average *χ*^2^ statistic of COAST was 37% higher than that of SKAT-O ALL and 4.8-times higher than that of SKAT-O PTV (**Supplemental Table 6**).

**Figure 5:**
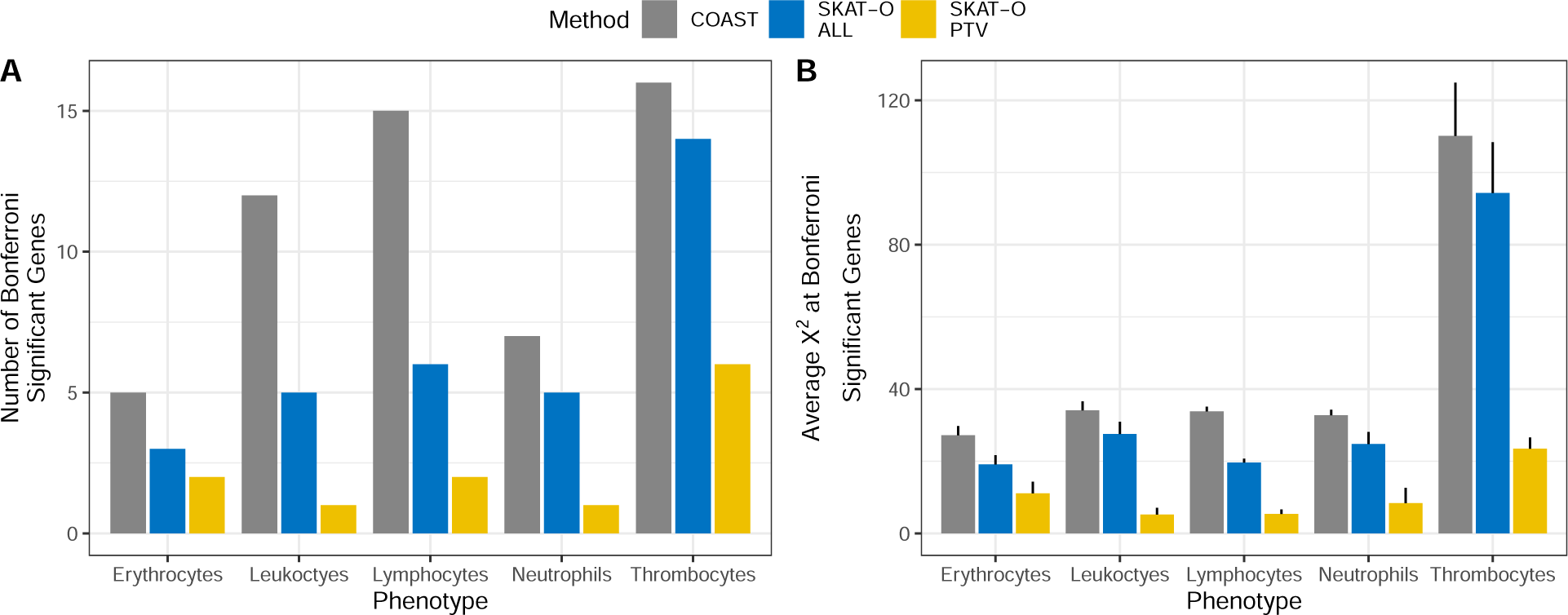
Number and significance of genes significantly associated with cell count phenotypes. **A.** Number of significantly associated genes by phenotype and association test. **B.** Average *χ*^2^ across the union of genes considered Bonferroni significant by any association test.

Among 61 total associations, 25 (41%) were unique to COAST, 33 (54%) were in common, and 3 (5%) were unique to SKAT-O ALL (**Supplemental Figure 13**). All associations except 1 (*TNXB* with thrombocytes, identified only by SKAT-O ALL) either had common variant support, from the GWAS catalog, rare variant support, from Genebass, or both (**Supplemental Tables 7-8**). One gene, *DOK2* (*p* = 4.8 × 10^−7^), associated with lymphocytes by COAST, did not previously reach significance in Genebass, although its association with missense variants was suggestive (*p* = 7.6 × 10^−6^). Many well-known gene-trait associations were recapitulated, including *HBB* (*p* = 1.2 × 10^−8^) for erythrocytes, *CXCR2* (2.6 × 10^−41^) for neutrophils, and *JAK2* (*p* = 2.2 × 10^−86^) for thrombocytes (**Figure 15**, Supplemental Figure 15).

For cell count traits, the pattern of effect sizes among genes identified by COAST was not always monotone increasing, although there were many such examples: *SH2B3* with erythrocytes, *S1PR2* with leukoctyes, *SBNO2* with lymphocotyes, and *JAK2* with thrombocytes (**Supplemental Figure 16**). Among genes identified as significant by SKAT-O ALL but not COAST, none had an allelic series pattern of effect sizes. **Figure 6** compares the average *χ*^2^ statistics of COAST and SKAT-O at empirical allelic series for both lipid and cell count traits. A gene was considered an empirical allelic series if it was significantly associated with the trait by any association test and the mean effect size increased monotonically from BMVs to DMVs to PTVs. Such genes are the intended targets of COAST, and COAST had the best power to uncover such genes.

**Figure 6:**
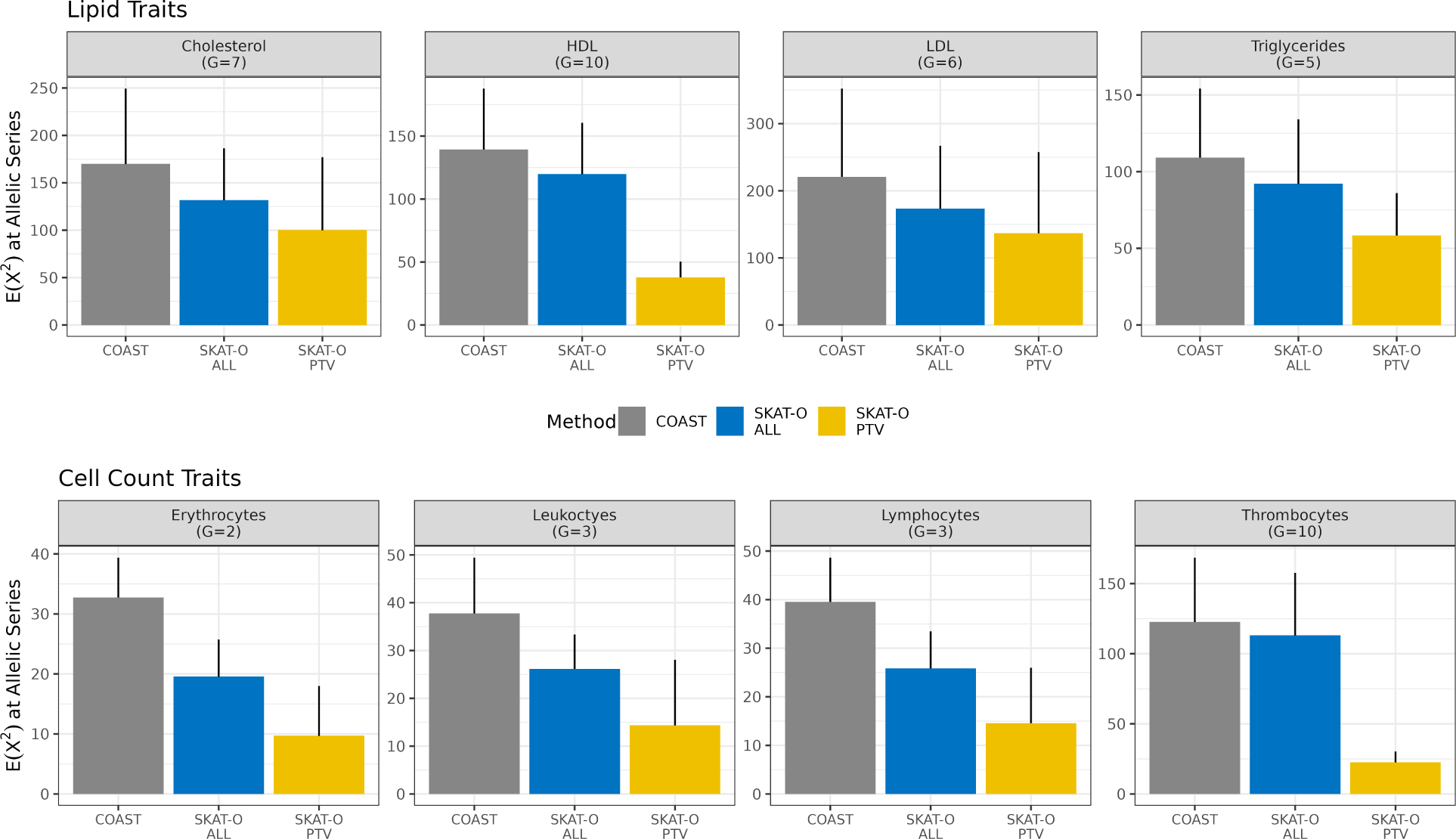
Average *χ*^2^ statistic at empirical allelic series. A gene was considered an empirical allelic series if it was significantly associated with the trait by any association test and the mean effect size increased monotonically from BMVs to DMVs to PTVs. Note that the mean effect size is subject to estimation error. The number of allelic series represented in each panel is denoted by *G*. No panel is shown for neutrophils because no genes with strictly monotonic effect sizes were identified.

## Discussion

We have developed a gene-based rare variant association test targeted to the identification of allelic series: genes where increasingly deleterious mutations have increasingly large phenotypic effects. While masks are widely used to select which variants to include in a rare variant analysis [20, 21], to our knowledge, COAST is the first association test that specifically seeks genes that harbor a dose-response relationship between mutational severity and phenotypic impact. In simulation studies, COAST controlled the type I error and tended to improve power when the pattern of effect sizes increased monotonically from BMVs to DMVs to PTVs. In applications to circulating lipid and cell count phenotypes from the UK Biobank, COAST consistently identified more Bonferroni significant associations than SKAT-O, and did so at a higher level of significance. In all cases, the additional genes detected by COAST had supporting evidence from at least one of Genebass or the GWAS catalog, mitigating the risk that these associations are spurious. Empirically, it was shown that in many cases the pattern of effect sizes for genes identified by COAST increased monotonically, and that for genes where the pattern of effect sizes was monotonic, COAST provided the greatest power.

COAST is not intended as a replacement for existing rare-variant associations tests, such as SKAT-O [16], ACAT-V [26], STAAR [19], or SAIGE-GENE+ [27]. These methods seek to identify the existence of a gene-trait association in general, placing no prior on what form that relationship should take. In contrast, COAST is tailored to the identification of genes that harbor a dose-response relationship. As such, the collection of genes targeted by COAST is only a subset of that targeted by existing methods. Empirically, the effect sizes of genes identified by COAST were not always monotone increasing in mutational severity. In part this is likely due to uncertainty in estimating the effect sizes of individual rare variants, however it is also due in part to COAST retaining power for detecting gene-trait associations that follow a partial allelic series pattern. In practice, when the goal of an analysis is to identify high-confidence allelic series, the initial results of COAST may require *post hoc* filtering to remove those genes whose effect size patterns are not monotone increasing.

COAST identified several gene-trait associations that are not present in Genebass but which garner support from the GWAS catalog: *ANGPTL4* (triglycerides), *A1CF* (triglycerides), and *DOK2* (lymphocytes). A low-frequency coding variant (rs116843064) in *ANGPTL4* was associated with HDL and triglycerides among 42K individuals of European ancestry (outside of UK Biobank) [41] and among 66K individuals of multiple ancestries (outside of the UK Biobank) [42]. A meta-analysis of 50K individuals associated rs116843064 with fasting triglyceride levels [43]. A candidate gene study associated rare inactivating mutations in *ANGPTL4* with reduced levels of fasting triglycerides and reduced odds of coronary artery disease [44]. Consistent with that work, 25 of the 27 rare variants in *ANGPTL4* considered in the present study, including all PTVs, were marginally associated with decreased triglyceride levels. A low-frequency coding variant (rs41274050) in *A1CF* was associated with triglycerides and total cholesterol in over 300K individuals of various ancestries, and the association was validated experimentally in a mouse model [45]. A large trans-ethnic analysis of 750K individuals, including UK Biobank, identified a low-frequency coding variant (rs56094005) in *DOK2* as associated with lymphocyte count, in addition to monocytes, neutrophils, and platelets [46, 47].

COAST has several limitations and opportunities for extension. The current implementation only incorporates rare coding variants annotated as a BMV, DMV, or PTV. While a clear ordering exists among the expected phenotypic impacts of these variant categories, in other cases the effect of a variant on gene function is complex and difficult to predict [48]. One possible extension of COAST is to assign a continuous measure of mutational severity, such as the CADD [49, 50], MACIE [51], or annotation principal component [19, 52] scores, to all variants within a gene, including non-coding variants. The continuous metric could then be cut into a finite number of intervals, and the variants categorized accordingly. These mutational severity intervals would replace the current categorization into BMV, DMV, and PTV. Open questions include what metric to use for measuring mutational severity and how to discretize it. Alternatively, the idea of categorizing could be discarded, and models could be defined that allow the expected value of the phenotype to vary continuous with, for example, the total value of the mutational severity metric aggregated across all rare variants within the gene.

We view the allelic series weights as hyperparameters that specify the pattern of effect sizes sought. The default values of (*w*_1_, *w*_2_, *w*_3_) = (1, 2, 3) for BMVs, DMVs, and PTVs specify a monotone increasing pattern. However, any choice of weights such that *w*_1_ ≤ *w*_2_ ≤ *w*_3_ suffices to tailor COAST for detecting allelic series. In general the optimal weighting scheme will depend on the genetic architecture of a given gene-trait pair. Nevertheless, a weighting scheme that is informed by the expected distribution of effect sizes for a particular phenotype would likely improve power. For example, data from a single large biobank might be split into training and evaluation sets. COAST with default weights could be run on the training data, and a collection of candidate allelic series identified. Among the candidate allelic series, the average effect sizes of BMVs, DMVs, and PTVs could be calculated. These average effect sizes would then serve as the weights (*w*_1_, *w*_2_, *w*_3_) when applying COAST to the evaluation data. Improving power by learning the allelic series weights empirically is a promising direction for future research.

Although COAST does not consider the effects of common variants, many prominent examples of allelic series have supporting evidence from both rare and common variants. For example, cholesterol levels were associated with common non-coding variants in a locus near *ANGPTL3* [53] and with rare loss-of-function variants in the same gene [54, 3]. Additionally, both a common missense variant and rare PTVs in *SLC30A8* were independently associated with type 2 diabetes [55]. Extending COAST to include common variant effects would increase power to detect allelic series that have both rare and common variant support.

Finally, a single gene may have a dose-response relationship with multiple related traits. For example, *APOB* was significantly associated with all of cholesterol, HDL, LDL, and triglycerides via COAST. In such cases, a multivariate outcome model, such as a linear mixed effects model, can improve power by simultaneously estimating the effects of rare variants on multiple correlated traits. Mixed effects modeling also enables the analysis of genetically related individuals, for example by including a random intercept with covariance proportional to the genetic relatedness matrix in the association model [56, 57]. Alternatively, COAST can be extended to non-quantitative phenotypes, such as binary, count, or time-to-event phenotypes, by generalizing the constituent models to generalized linear models or Cox proportional hazards models. These extensions are under active development.

## Acknowledgements

The authors would like to acknowledge Baris Ungun for contributions to curating the UK Biobank data, and James Warren for contributions to analysis infrastructure, and to thank the participants of the UK Biobank, whose data were used with permission.

## Author Contributions

TWS, DK, and FPC conceived of the project. FPC curated the phenotypic data. TWS curated the genetic data and performed the proof-of-principle analysis. ZRM developed the method and performed the final analysis. All authors provided scientific input. CK prepared the software for production. ZRM and TWS wrote the first draft of the manuscript. All authors contributed to critical revision of the final manuscript.

## Data and Code Availability

This work uses genotypes and phenotypes from the UK Biobank (https://www.ukbiobank.ac.uk/) accessed pursuant to approved application number **51766**. The coding-variant allelic series test has been implemented as an R [58] package, which is available on GitHub at https://github.com/insitro/AllelicSeries.

## References

[1] B McClintock. The relation of homozygous deficiencies to mutations and allelic series in maize. Genetics, 29(5):478–502, 1944.

[2] PM Plenge, EM Scolnick, and D Altshuler. Validating therapeutic targets through human genetics. Nat Rev Drug Discov, 12(8):581–594, 2013.

[3] K Musunuru and S Kathiresan. Genetics of common, complex coronary artery disease. Cell, 177(1):132–145, 2019.

[4] CA Dendrou, A Cortes, L Shipman, et al. Resolving tyk2 locus genotype-to-phenotype differences in autoimmunity. Sci Transl Med, 8(363):363ra149, 2016.

[5] SM Hoy. Deucravacitinib: First approval. Drugs, 82(17):1671–1679, 2022.

[6] C Sudlow, J Gallacher, N Allen, et al. Uk biobank: an open access resource for identifying the causes of a wide range of complex diseases of middle and old age. PLoS Med, 12(3):e1001779, 2015.

[7] PM Visscher, NR Wray, Q Zhang, P Sklar, MI McCarthy, MA Brown, and J Yang. 10 years of gwas discovery: Biology, function, and translation. Am J Hum Genet, 101(1):5–22, 2017.

[8] PL Auer and G Lettre. Rare variant association studies: considerations, challenges and opportunities. Genome Med, 7(1):16, 2015.

[9] JA Kosmicki, CL Churchhouse, MA Rivas, and BN Neale. Discovery of rare variants for complex phenotypes. Hum Genet, 135(6):625–634, 2016.

[10] S Lee, GA Abecasis, M Boehnke, and X Lin. Rare-variant association analysis: Study designs and statistical tests. Am J Hum Genet, 95(1):5–23, 2014.

[11] J Asimit and E Zeggini. Rare variant association analysis methods for complex traits. Annu Rev Genet, 44:293–308, 2010.

[12] BE Madsen and SR Browning. A groupwise association test for rare mutations using a weighted sum statistic. PLoS Genet, 5:e1000384, 2009.

[13] AP Morris and E Zeggini. An evaluation of statistical approaches to rare variant analysis in genetic association studies. Genet Epidemiol, 34(2):188–193, 2010.

[14] S Morgenthaler and WG Thilly. A strategy to discover genes that carry multi-allelic or mono-allelic risk for common diseases: a cohort allelic sums test (cast). Mutat Res, 615:28–56, 2014.

[15] MC Wu, S Lee, T Cai, Y Li, M Boehnke, and X Lin. Rare-variant association testing for sequencing data with the sequence kernel association test. Am J Hum Genet, 89(1):82–93, 2011.

[16] Seunggeun Lee, Mary J. Emond, Michael J. Bamshad, Kathleen C. Barnes, Mark J. Rieder, Deborah A. Nickerson, David C. Christiani, Mark M. Wurfel, and Xihong Lin. Optimal unified approach for rare-variant association testing with application to small-sample case-control whole-exome sequencing studies. The American Journal of Human Genetics, 91(2):224–237, 2012.

[17] Z He, B Xu, S Lee, and I Ionita-Laza. Unified sequence-based association tests allowing for multiple functional annotations and meta-analysis of noncoding variation in metabochip data. Am J Hum Genet, 101(3):340–352, 2017.

[18] Y Ma and P Wei. Funspu: A versatile and adaptive multiple functional annotation-based association test of whole-genome sequencing data. PLoS Genet, 15(4):e1008081, 2019.

[19] X Li, Z Li, H Zhou, et al. Dynamic incorporation of multiple in silico functional annotations empowers rare variant association analysis of large whole-genome sequencing studies at scale. Nat Genet, 52(9):969–983, 2020.

[20] J Mbatchou, L Barnard, J Backman, et al. Computationally efficient whole-genome regression for quantitative and binary traits. Nat Genet, 53(7):1097–1103, 2021.

[21] Z Li, X Li, H Zhou, et al. A framework for detecting noncoding rare-variant associations of large-scale whole-genome sequencing studies. Nat Methods, 19(12):1599–1611, 2022.

[22] W McLaren, L Gil, SE Hunt, et al. The ensembl variant effect predictor. Genome Biol, 17(1):122, 2016.

[23] KJ Karczewski, M Solomonson, KR Chao, et al. Systematic single-variant and gene-based association testing of thousands of phenotypes in 394,841 uk biobank exomes. Cell Genomics, 2(9):100168, 2022.

[24] G Seber. The Linear Model and Hypothesis. Springer, 2015.

[25] Y Liu and J Xie. Cauchy combination test: a powerful test with analytic p-value calculation under arbitrary dependency structures. Journal of the American Statistical Association, 115(529):393–402, 2020.

[26] Y Liu, S Chen, Z Li, AC Morrison, E Boerwinkle, and X Lin. Acat: A fast and powerful p value combination method for rare-variant analysis in sequencing studies. Am J Hum Genet, 104(3):410–421, 2019.

[27] W Zhou, W Bi, Z Zhao, et al. Saige-gene+ improves the efficiency and accuracy of set-based rare variant association tests. Nat Genet, 54(10):1466–1469, 2022.

[28] HM Amemiya, A Kundaje, and AP Boyle. The encode blacklist: Identification of problematic regions of the genome. Sci Reports, 9:9354, 2019.

[29] H Li. Toward better understanding of artifacts in variant calling from high-coverage samples. Bioinformatics, 30(20):2843–2851, 2014.

[30] Adzhubei IA, Schmidt S, Peshkin L, et al. A method and server for predicting damaging missense mutations. Nat Methods, 7(4):248–249, 2010.

[31] Ng PC and Henikoff S. Predicting deleterious amino acid substitutions. Genome Res, 11(5):863–874, 2001.

[32] ET Cirulli, S White, RW Read, et al. Genome-wide rare variant analysis for thousands of phenotypes in over 70,000 exomes from two cohorts. Nat Commun, 11:542, 2020.

[33] CV Van Hout, I Tachmazidou, JD Backman, et al. Exome sequencing and characterization of 49,960 individuals in the uk biobank. Nature, 586:749–756, 2020.

[34] KJ Karczewski, LC Francioli, G Tiao, et al. The mutational constraint spectrum quantified from variation in 141,456 humans. Nature, 581:434–443, 2020.

[35] C Bycroft, C Freeman, D Petkova, et al. The uk biobank resource with deep phenotyping and genomic data. Nature, 562:203–209, 2018.

[36] Hail team. hail 0.2. https://github.com/hail-is/hail.

[37] ZR McCaw, JM Lane, R Saxena, S Redline, and X Lin. Operating characteristics of the rank-based inverse normal transformation for quantitative trait analysis in genome-wide association studies. Biometrics, 76(4):1262–1272, 2020.

[38] A Buniello, JAL MacArthur, M Cerezo, et al. The nhgri-ebi gwas catalog of published genome-wide association studies, targeted arrays and summary statistics 2019. Nucleic Acids Research, 47(D1):D1005–D1012, 2019.

[39] Gwas catalog: The NHGRI-EBI catalog of human genome-wide association studies. https://www.ebi.ac.uk/gwas/. Accessed: 2022-11-23.

[40] Genebass: gene-based association summary statistics. https://app.genebass.org/. Accessed: 2022-11-23.

[41] GM Peloso, PL Auer, JC Bis, et al. Association of low-frequency and rare coding-sequence variants with blood lipids and coronary heart disease in 56,000 whites and blacks. Am J Hum Genet, 94(2):223–232, 2014.

[42] MS Selvaraj, X Li, Z Li, et al. Whole genome sequence analysis of blood lipid levels in >66,000 individuals. Nat Commun, 13(1):5995, 2022.

[43] EM van Leeuwen, A Sabo, JC Bis, et al. Meta-analysis of 49,549 individuals imputed with the 1000 genomes project reveals an exonic damaging variant in angptl4 determining fasting tg levels. J Med Genet, 53(7):441–449, 2016.

[44] FE Dewey, V Gusarova, C O’Dushlaine, et al. Inactivating variants in angptl4 and risk of coronary artery disease. N Engl J Med, 374(12):1123–33, 2016.

[45] DJ Liu, GM Peloso, H Yu, et al. Exome-wide association study of plasma lipids in >300,000 individuals. Nat Genet, 49(12):1758–1766, 2017.

[46] MH Chen, LM Raffield, A Mousas, et al. Trans-ethnic and ancestry-specific blood-cell genetics in 746,667 individuals from 5 global populations. Cell, 182(5):1198–1213, 2020.

[47] D Vuckovic, EL Bao, P Akbari, et al. The polygenic and monogenic basis of blood traits and diseases. Cell, 182(5):1214–1231, 2020.

[48] T Lappalainen and DG MacArthur. From variant to function in human disease genetics. Science, 24(373):1464–1468, 2021.

[49] M Kircher, DM Witten, P Jain, et al. A general framework for estimating the relative pathogenicity of human genetic variants. Nat Genet, 46(3):310–315, 2014.

[50] P Rentzsch, DM Witten, GM Cooper, J Shendure, and M Kircher. Cadd: predicting the deleteriousness of variants throughout the human genome. Nucleic Acids Research, 47(D1):D886–D894, 2019.

[51] X Li, G Yung, H Zhou, et al. A multi-dimensional integrative scoring framework for predicting functional variants in the human genome. Am J Hum Genet, 109(3):446–456, 2022.

[52] H Zhou, T Arapoglou, X Li, et al. Favor: functional annotation of variants online resource and annotator for variation across the human genome. Nucleic Acids Res, 51(D1):D1300–D1311, 2023.

[53] T Teslovich, K Musunuru, A Smith, et al. Biological, clinical and population relevance of 95 loci for blood lipids. Nature, 466:707–713, 2010.

[54] FE Dewey, V Gusarova, RL Dunbar, et al. Genetic and pharmacologic inactivation of angptl3 and cardiovascular disease. N Engl J Med, 377(3):211–221, 2017.

[55] J Flannick, JM Mercader, C Fuchsberger, et al. Exome sequencing of 20,791 cases of type 2 diabetes and 24,440 controls. Nature, 570:71–76, 2019.

[56] PR Loh, G Tucker, BK Bulik-Sullivan, et al. Efficient bayesian mixed-model analysis increases association power in large cohorts. Nat Genet, 47(3):284–290, 2015.

[57] H Chen, C Wang, MP Conomos, et al. Control for population structure and relatedness for binary traits in genetic association studies via logistic mixed models. Am J Hum Genet, 98(4):653–666, 2016.

[58] R Core Team. R: A Language and Environment for Statistical Computing. R Foundation for Statistical Computing, Vienna, Austria, 2022.

