## Supplemental for "An allelic series rare variant association test for candidate gene discovery"

April 28, 2023

[1] Insitro, South San Francisco, CA, USA.

[2] Department of Informatics, Technical University of Munich, Munich, Germany.

### 1 Supplemental Methods

#### 1.1 Standard Wald Test

Consider the model:

$$\boldsymbol{y} = \boldsymbol{G}\boldsymbol{\beta} + \boldsymbol{X}\boldsymbol{\gamma} + \boldsymbol{\epsilon}, \tag{1}$$

where $\boldsymbol{y}$ is an $N \times 1$ phenotype vector, $\boldsymbol{G}$ is an $N \times P$ genotype matrix, $\boldsymbol{X}$ is an $N \times Q$ covariate matrix, and $\boldsymbol{\epsilon}$ is an $N \times 1$ residual with mean $\mathbb{E}(\boldsymbol{\epsilon}|\boldsymbol{G}, \boldsymbol{X}) = \boldsymbol{0}$ and variance $\mathbb{V}(\boldsymbol{\epsilon}|\boldsymbol{G}, \boldsymbol{X}) = \sigma^2 \boldsymbol{I}$. For convenience, the intercept has been incorporated into $\boldsymbol{X}$. With different choices of $\boldsymbol{G}$, (1) encompasses all of the allelic series burden models:

- For the baseline model, $\boldsymbol{G}$ is a $N \times 3$ matrix, where the 3 columns are the counts $(N_1, N_2, N_3)$ of BMVs, DMVs, and PTVs respectively.
- For the allelic series sum model, $\boldsymbol{G}$ is an $N \times 1$ matrix, with $G_i = \sum_{l=1}^{3} w_l N_{il}$.
- For the allelic series max model, $\boldsymbol{G}$ is an $N \times 1$ matrix, with $G_i = \max_{l=1}^{3} w_l N_{il}$.

Let $\hat{\boldsymbol{\beta}}$ denote the ordinary least squares (OLS) estimator of $\boldsymbol{\beta}$, which is expressible as:

$$\hat{\boldsymbol{\beta}} = (\boldsymbol{G}^T \boldsymbol{P}_X^{\perp} \boldsymbol{G})^{-1} \boldsymbol{G}^T \boldsymbol{P}_X^{\perp} \boldsymbol{y},$$

where $\boldsymbol{P}_X^{\perp} = \boldsymbol{I} - \boldsymbol{X}(\boldsymbol{X}^T\boldsymbol{X})^{-1}\boldsymbol{X}^T$. The standard Wald test of $H_0 : \boldsymbol{\beta} = \boldsymbol{0}$ is:

$$T_{\text{Wald}} = \hat{\boldsymbol{\beta}}^T \left\{ \mathbb{V}(\hat{\boldsymbol{\beta}}|\boldsymbol{G}, \boldsymbol{X}) \right\}^{-1} \hat{\boldsymbol{\beta}}. \tag{2}$$

Here $\mathbb{V}(\hat{\boldsymbol{\beta}})$ is the sampling variability of the OLS estimator:

$$\mathbb{V}(\hat{\boldsymbol{\beta}}|\boldsymbol{G}, \boldsymbol{X}) = \sigma^2 (\boldsymbol{G}^T \boldsymbol{P}_X^{\perp} \boldsymbol{G})^{-1},$$

and $\sigma^2$ is estimated by:

$$\hat{\sigma}^2 = (N - P - Q)^{-1} \boldsymbol{y}^T \boldsymbol{P}_{XG}^{\perp} \boldsymbol{y}, \tag{3}$$

where:

$$\boldsymbol{P}_{XG}^{\perp} = \boldsymbol{I} - \boldsymbol{Z}(\boldsymbol{Z}^T\boldsymbol{Z})^{-1}\boldsymbol{Z}^T, \qquad \boldsymbol{Z} = (\boldsymbol{G}, \boldsymbol{X}).$$

Under $H_0$, $T_{\text{Wald}} \sim \chi_{\nu}^2(0)$ where the degrees of freedom $\nu$ is the number of columns in $\boldsymbol{G}$.

#### 1.2 Standard Score Test

The score for evaluating $H_0 : \boldsymbol{\beta} = \boldsymbol{0}$ is:

$$\boldsymbol{S} = \boldsymbol{G}^T\boldsymbol{P}_X^{\perp}\boldsymbol{y}.$$

The variance of the score is:

$$\mathbb{V}(\boldsymbol{S}|\boldsymbol{G}, \boldsymbol{X}) = \sigma^2\boldsymbol{G}^T\boldsymbol{P}_X^{\perp}\boldsymbol{G},$$

where $\sigma^2$ is estimated by:

$$\tilde{\sigma}^2 = (N - Q)^{-1}\boldsymbol{y}^T\boldsymbol{P}_X^{\perp}\boldsymbol{y}. \tag{4}$$

Overall, the standard score statistic is:

$$T_{\text{Score}} = \boldsymbol{S}^T\left\{\mathbb{V}(\boldsymbol{S}|\boldsymbol{G}, \boldsymbol{X})\right\}^{-1}\boldsymbol{S}. \tag{5}$$

Like the Wald statistic, under $H_0$, $T_{\text{Score}} \sim \chi_{\nu}^2(0)$. Multiplying out (2) and (5) shows that both take the form:

$$T = \frac{\boldsymbol{y}^T\boldsymbol{P}_X^{\perp}\boldsymbol{G}(\boldsymbol{G}^T\boldsymbol{P}_X^{\perp}\boldsymbol{G})^{-1}\boldsymbol{G}^T\boldsymbol{P}_X^{\perp}\boldsymbol{y}}{\sigma^2},$$

the only difference being that the Wald test estimates $\sigma^2$ by (3), which requires $\boldsymbol{G}$, whereas the score test estimates $\sigma^2$ by (4), which does not require $\boldsymbol{G}$.

### 2 Supplemental Results

#### 2.1 Simulation Studies

##### 2.1.1 Type I Error

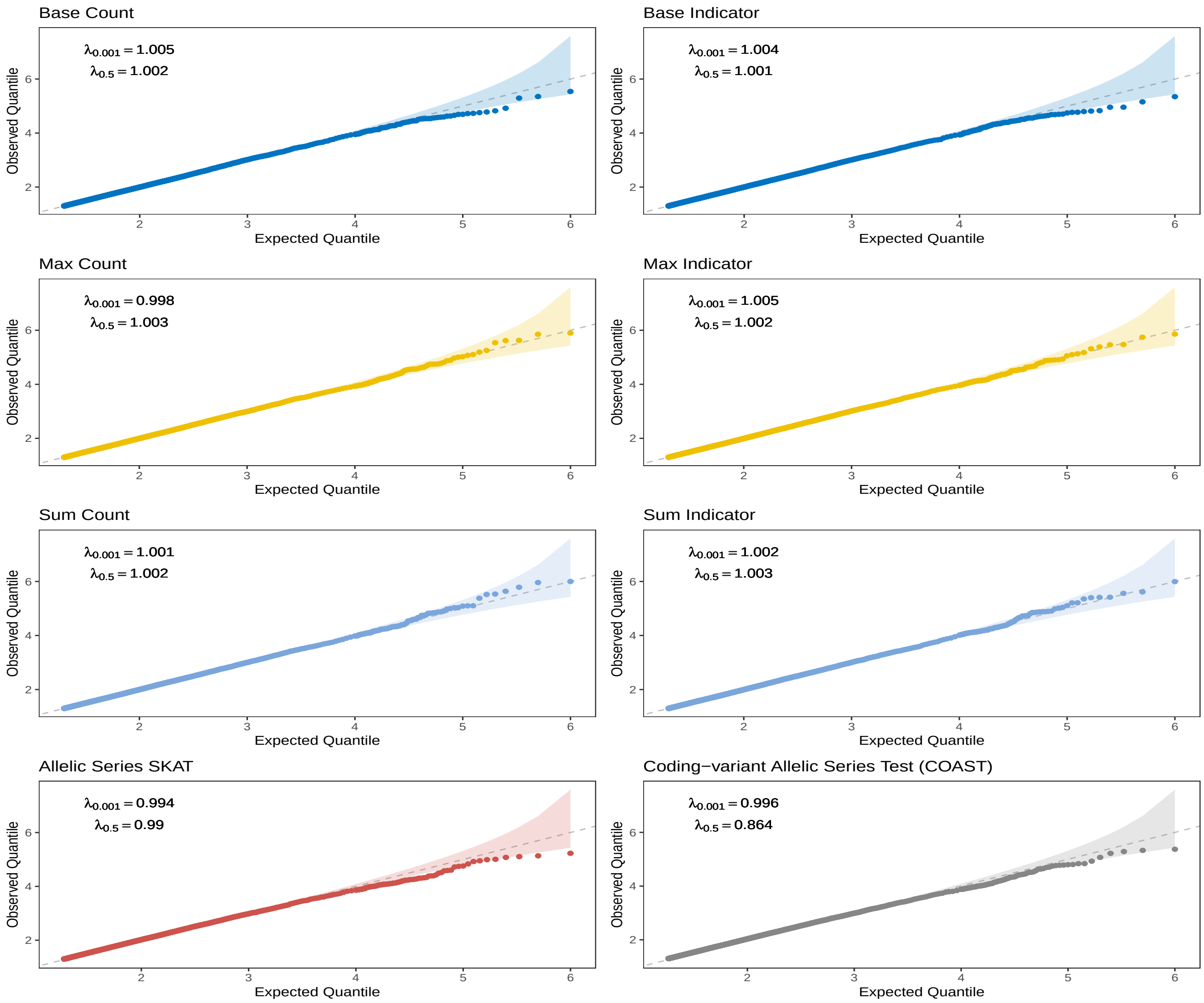

Figure 1: **Uniform quantile-quantile plots for the allelic series component tests under the null hypothesis of no association, using real genotypes extracted from the APOB gene.** Results are aggregated across $10^6$ simulations at sample size 10K. $\lambda_p$ is the genomic inflation factor calculated at the $p$th percentile. Adherence to the dashed 45° line indicates that the observed p-values are uniformly distributed.

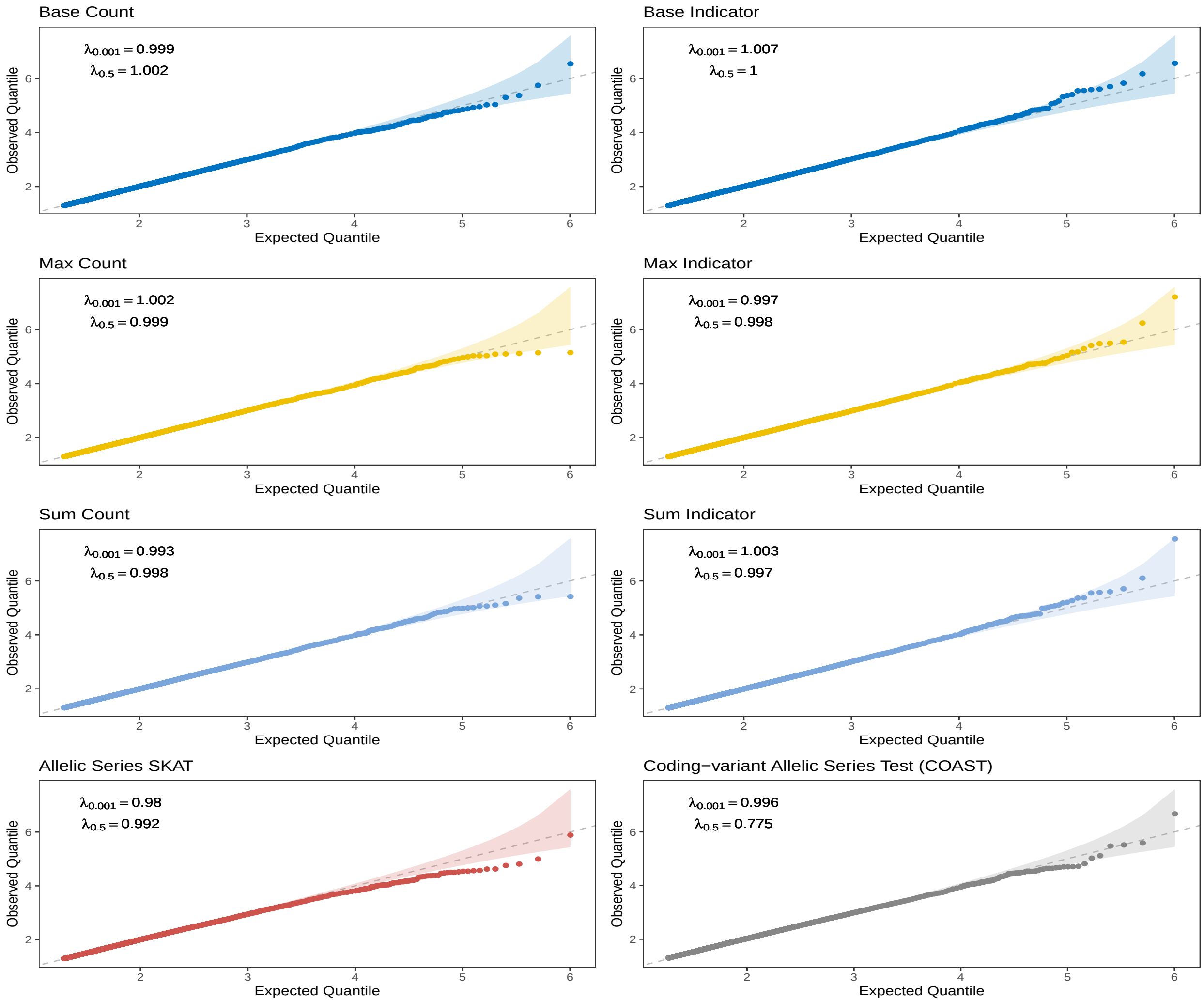

Figure 2: **Uniform quantile-quantile plots for the allelic series component tests under the null hypothesis of no association, using genotypes in linkage equilibrium.** Results are aggregated across $10^6$ simulations at sample size 10K. $\lambda_p$ is the genomic inflation factor calculated at the $p$th percentile. Adherence to the dashed 45° line indicates that the observed p-values are uniformly distributed.

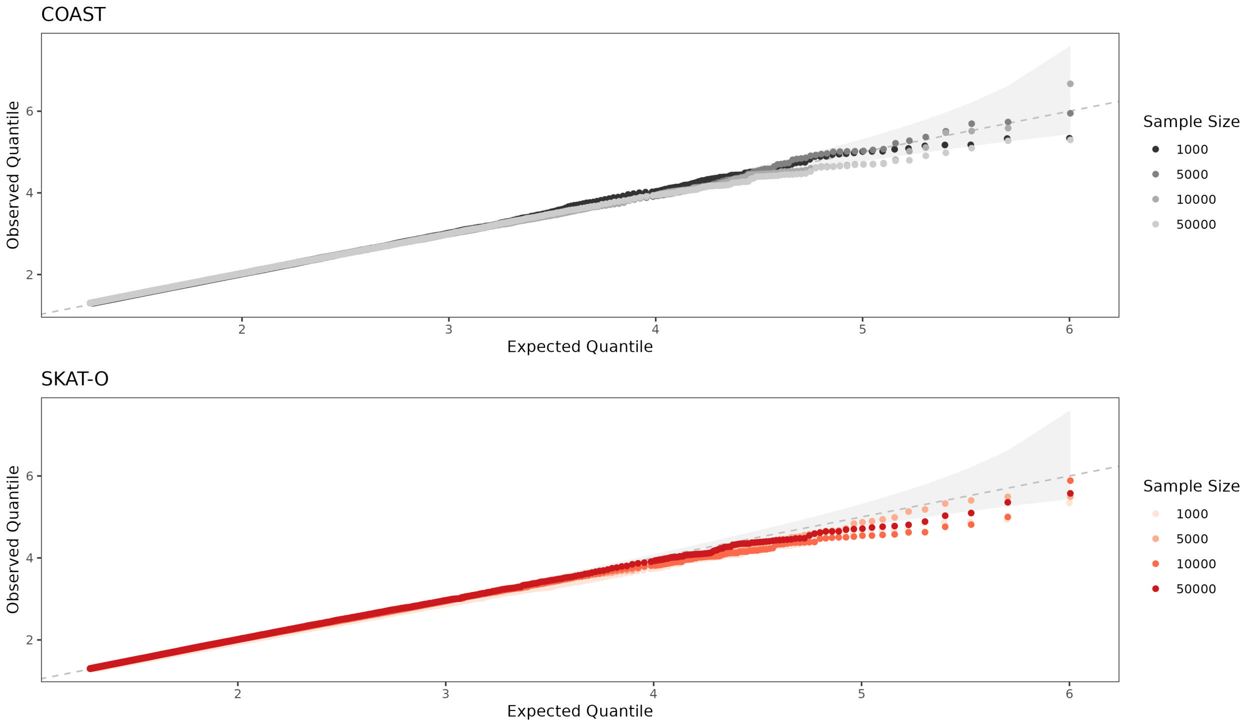

Figure 3: **Uniform quantile-quantile plots for the allelic series omnibus test (COAST) and SKAT-O under the null hypothesis of no association, using genotypes in linkage equilibrium.** Results are aggregated across $10^6$ simulations at sample sizes between 1K and 50K. Adherence to the dashed 45° line indicates that the observed p-values are uniformly distributed.

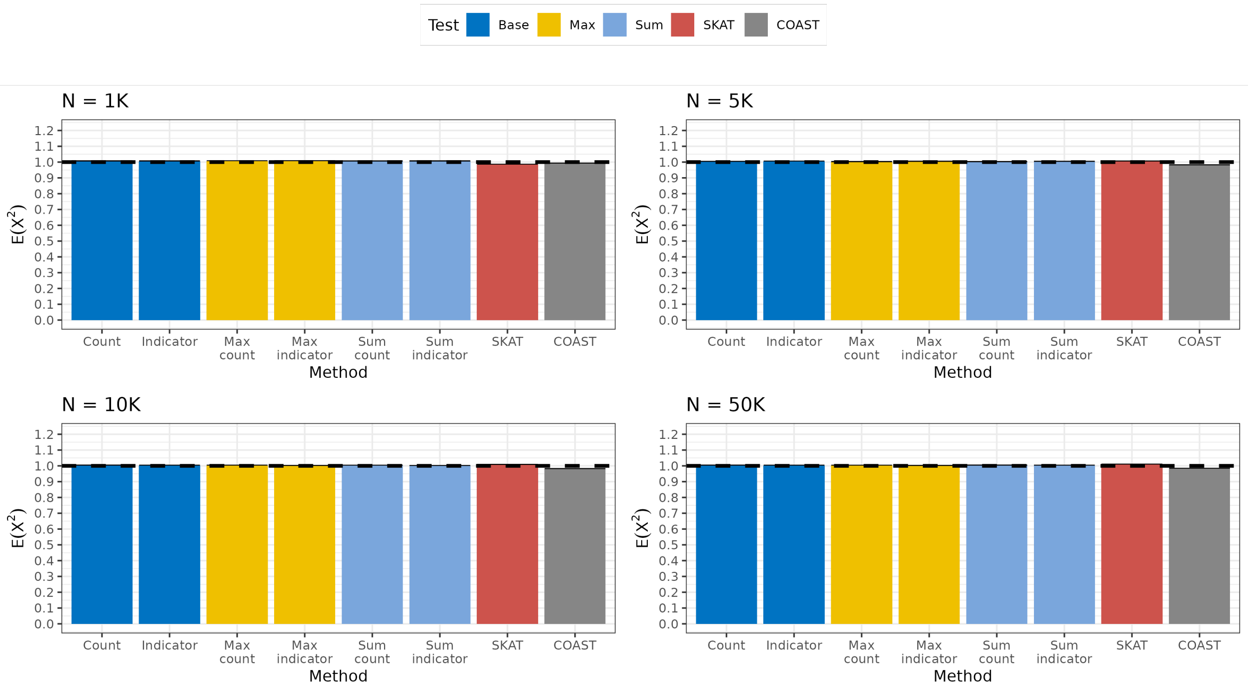

Figure 4: **Expected $\chi^2$ statistics of the allelic series component tests under the null hypothesis of no association.** Results are aggregated across $10^6$ simulations at sample sizes between 1K and 50K. Expected $\chi^2$ statistics of the allelic series test components as well as the overall omnibus test. A value of 1.0 is expected under the null.

##### 2.1.2 Power

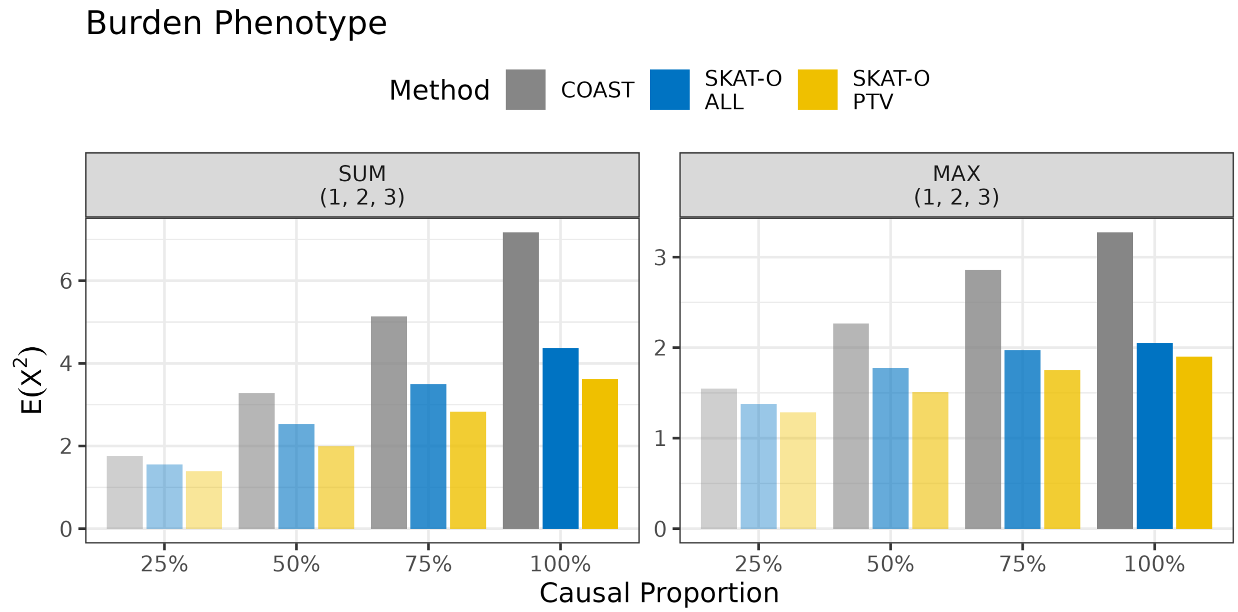

Figure 5: **Power to detect an allelic series under the SUM and MAX burden architectures.** The sample size is $N = 10^4$. Genes included $10^2$ variants with a BMV:DMV:PTV ratio of 5:4:1. Results are aggregated across $10^4$ simulations. The tuple in the panel heading denotes the relative effect sizes $(\beta_1, \beta_2, \beta_3)$ of benign missense variants, deleterious missense variants, and protein truncating variants. The allelic series weights are $w = (1, 2, 3)$. Note that the SUM and MAX architectures are incompatible with the SKAT phenotype.

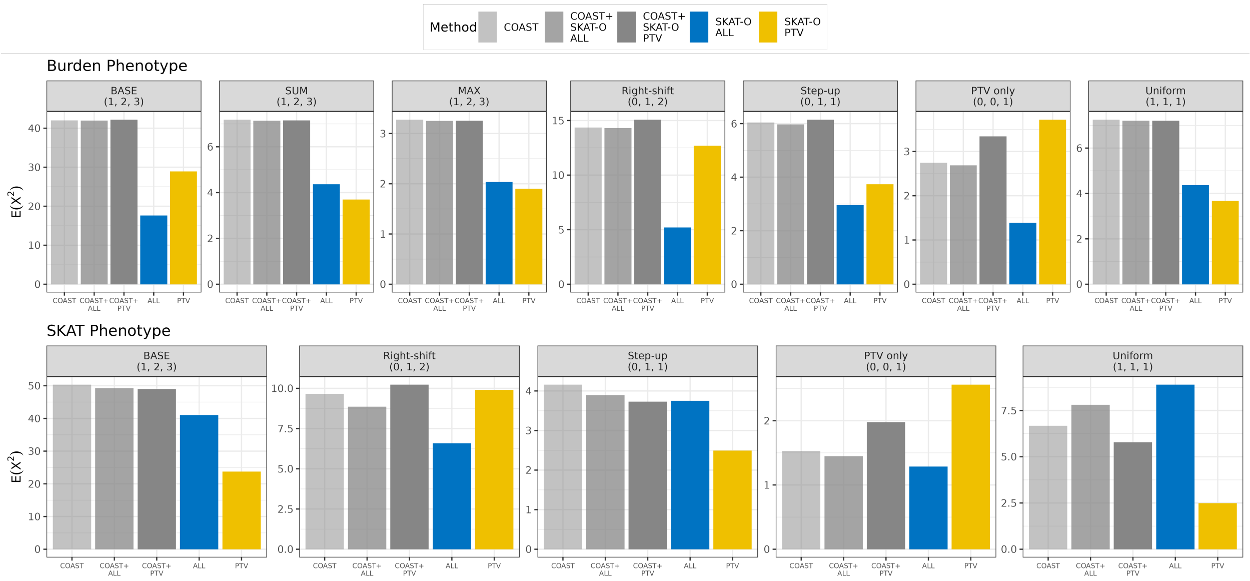

Figure 6: **Power to detect an allelic series under various genetic architectures.** The sample size is $N = 10^4$, each gene contained $10^2$ variants, and results are aggregated across $10^4$ simulations. Each tick corresponds to a different genetic architecture, including the BASE, SUM, and MAX architectures. The tuple above each panel denotes the relative effect sizes $(\beta_1, \beta_2, \beta_3)$ of benign missense variants, deleterious missense variants, and protein truncating variants. The allelic series weights remained fixed at $w = (1, 2, 3)$. For the burden phenotype, the effect sizes of all variants were in the same direction, whereas for the SKAT phenotype, the magnitude and direction of effect were randomized. The SUM and MAX models are omitted for the SKAT phenotype because these architectures are incompatible with randomized signs.

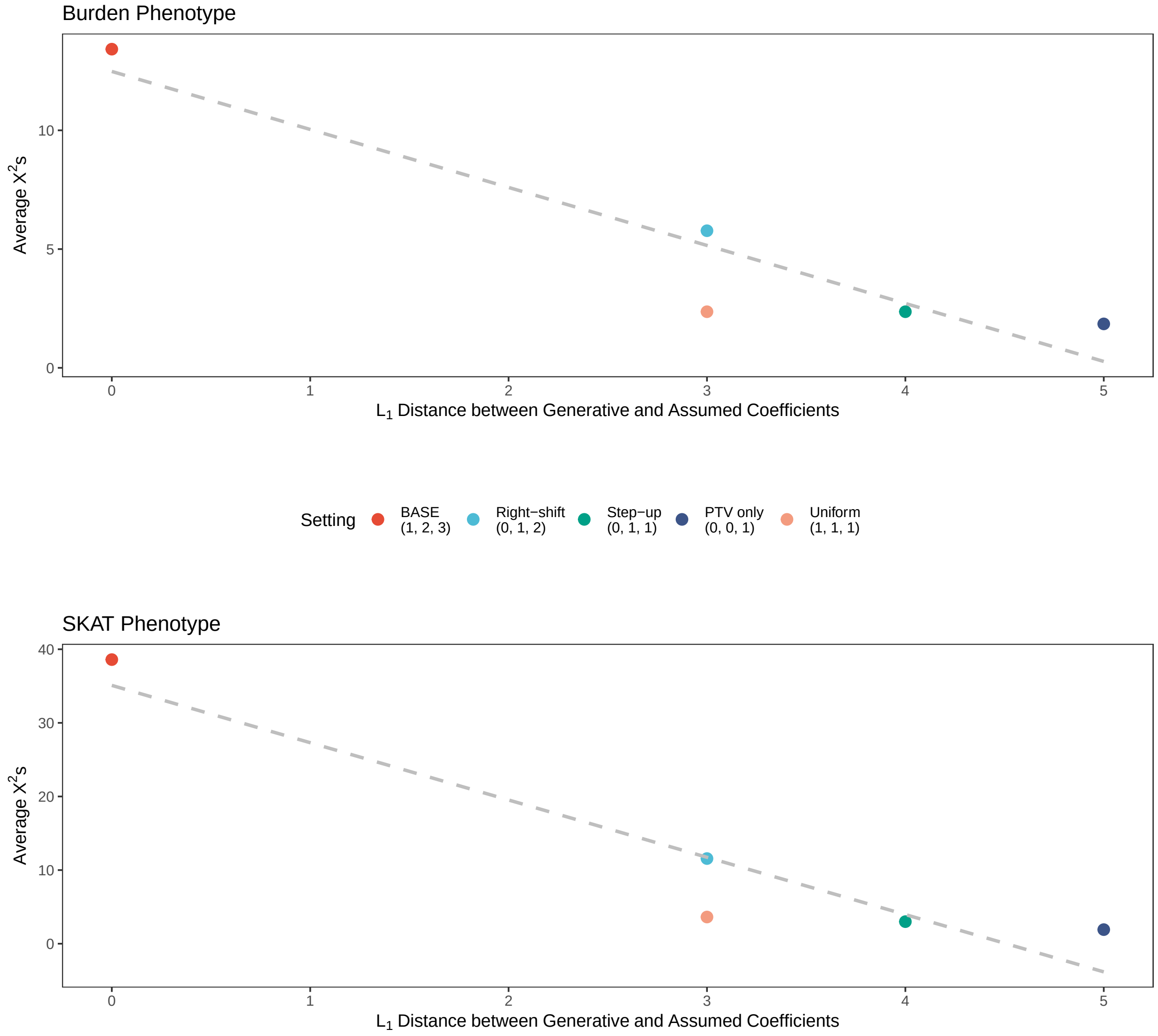

Figure 7: **Power of the coding-variant allelic series test by $L_1$ distance of the generative model from that assumed by COAST.** The sample size is $N = 10^3$, each gene contained $10^2$ variants, and results are aggregated across $10^4$ simulations. Color encodes the genetic architecture, with the tuple encoding the relative effect sizes $(\beta_1, \beta_2, \beta_3)$ of benign missense variants, deleterious missense variants, and PTVs. The dashed line is the linear trend. COAST assumes $(\beta_1, \beta_2, \beta_3) \propto (1, 2, 3)$. For the burden phenotype, the effect sizes of all variants were in the same direction, whereas for the SKAT phenotype, the direction of effect was randomized. The SUM and MAX models they are not nested within the baseline allelic series model (and therefore do not have comparable $\chi^2$ statistics).

#### 2.2 Circulating Lipid Phenotypes

##### 2.2.1 Figures

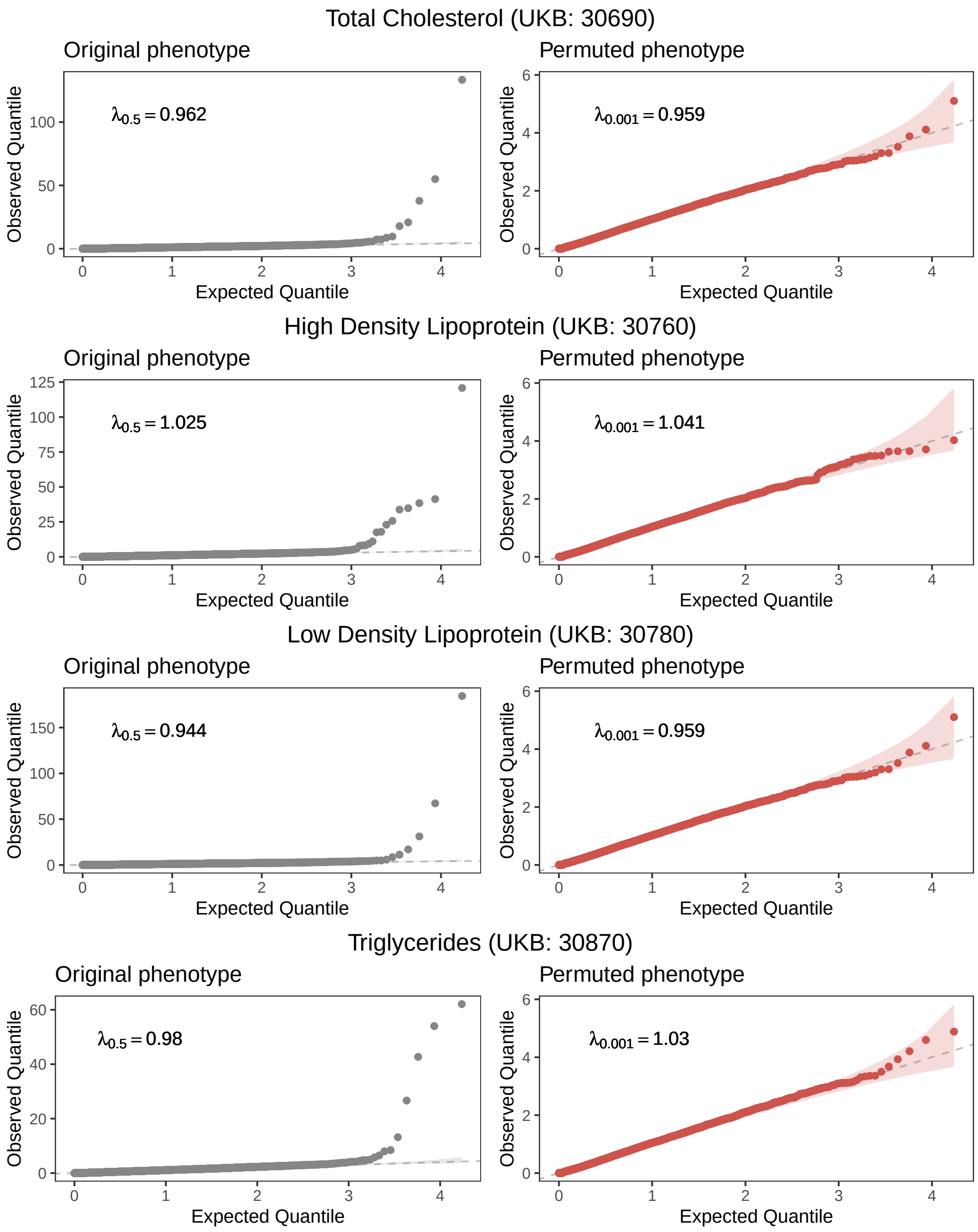

Figure 8: **Uniform quantile-quantile plots for COAST p-values on observed and permuted lipid phenotypes.** $\lambda_p$ is the genomic inflation factor calculated at the $p$th percentile. Note the difference in scale of the Y-axis between the original (left) and permuted phenotype (right).

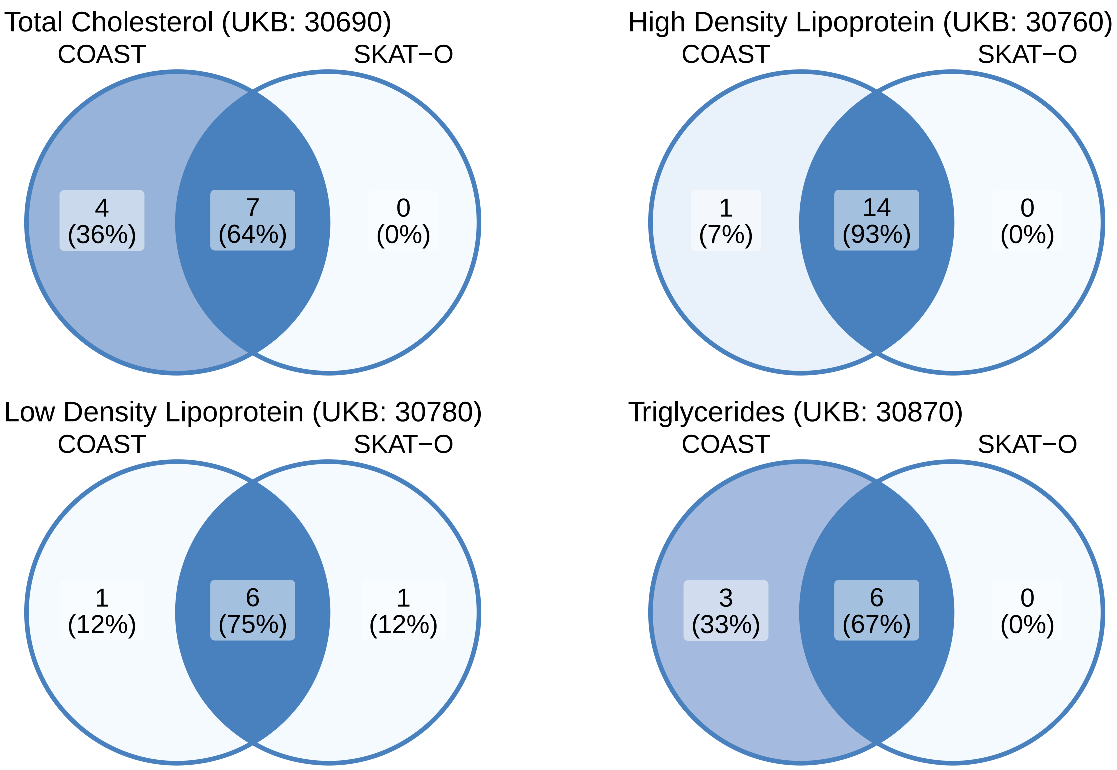

Figure 9: **Venn diagrams for the numbers of Bonferroni significant genes by association test for circulating lipid phenotypes.** COAST is the coding-variant allelic series test. SKAT-O is applied to all rare coding variants.

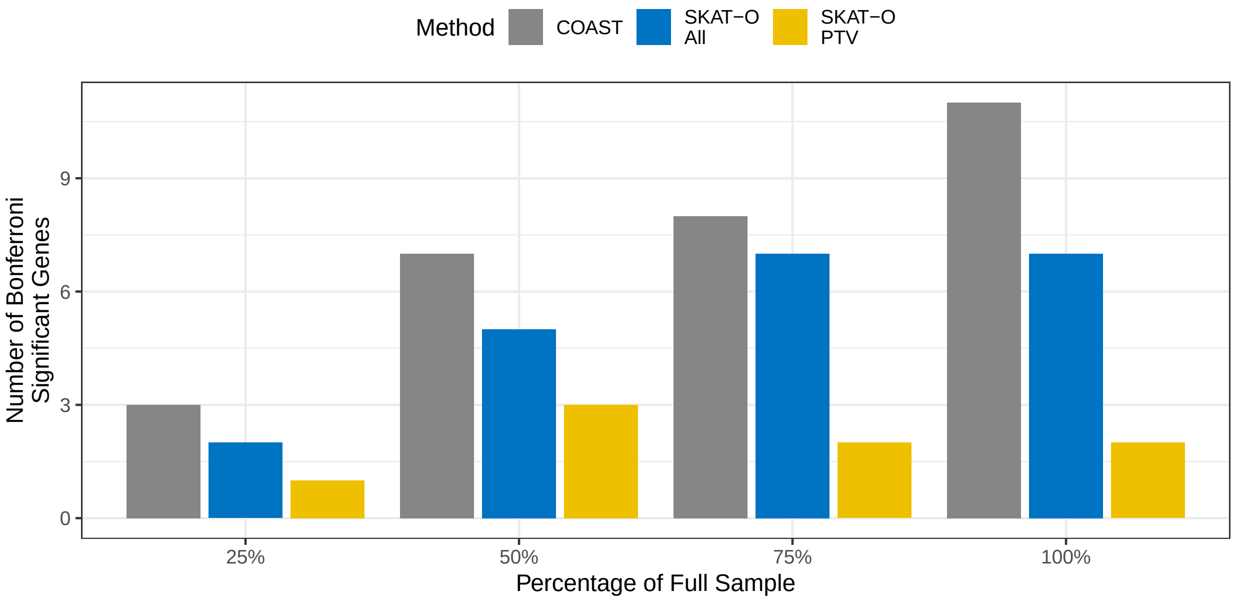

Figure 10: **Number of genes significantly associated with circulating cholesterol, by association test, as a function of the available sample size.** The total sample of size 145,735 was down-sampled to 75%, 50%, and 25% of available subjects. The samples were nested such that all subjects in a smaller sample were included in all larger samples.

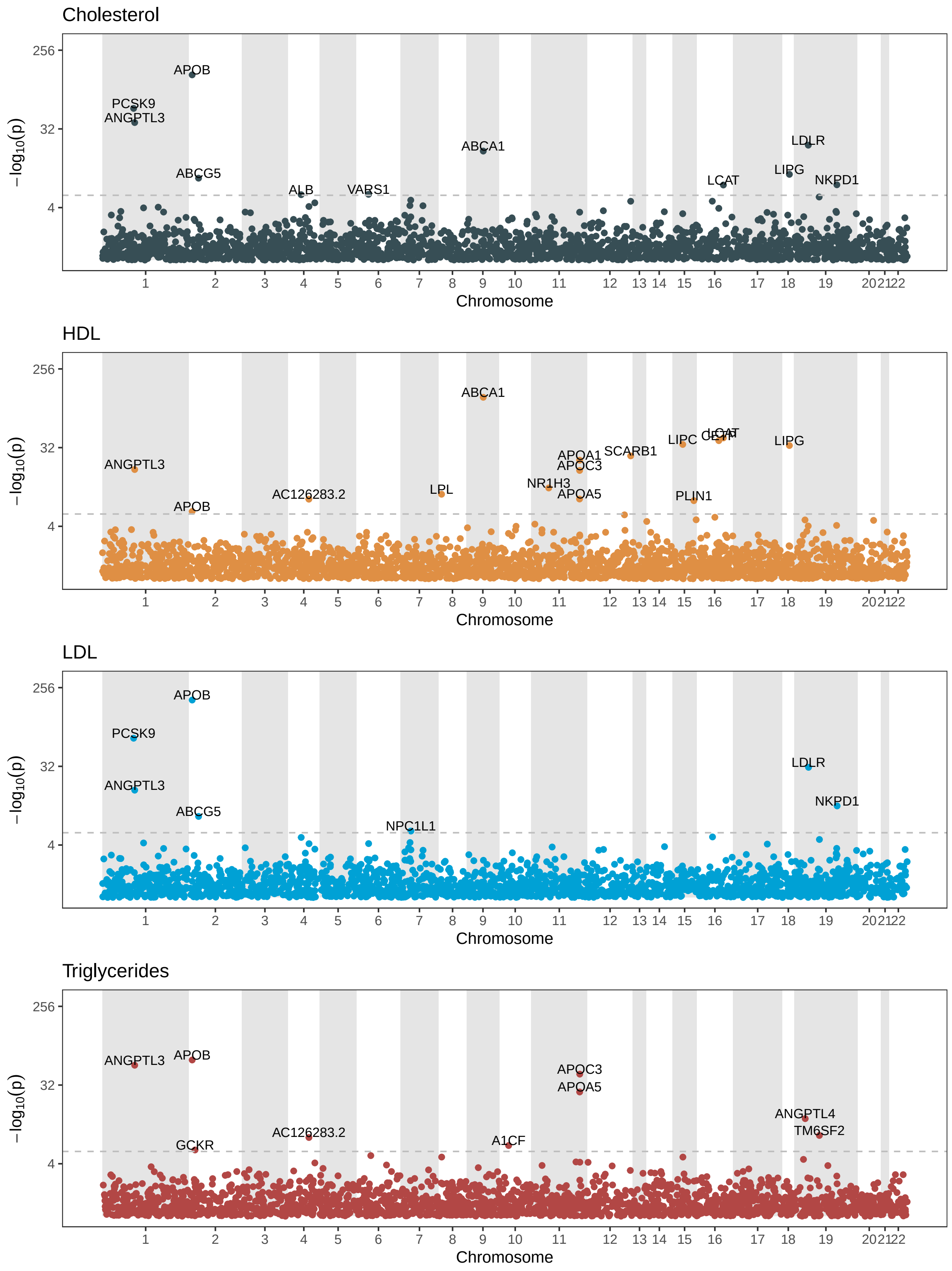

Figure 11: **Stratified Manhattan plots for allelic series (COAST) analysis of circulating lipid phenotypes.**

##### 2.2.2 Tables

Table 1: **Counts of Bonferroni significant genes by association test for circulating lipid phenotypes.** COAST is the coding-variant allelic series test, ALL is SKAT-O applied to all variants, and PTV is SKAT-O applied to protein truncating variants. COAST/ALL is the ratio of COAST to ALL and COAST/PTV is the ratio of COAST to PTV.

| Trait | COAST | ALL | PTV | COAST/ALL | COAST/PTV |
|---|---|---|---|---|---|
| Cholesterol | 11.00 | 7.00 | 2.00 | 1.57 | 5.50 |
| HDL | 15.00 | 14.00 | 6.00 | 1.07 | 2.50 |
| LDL | 7.00 | 7.00 | 2.00 | 1.00 | 3.50 |
| Triglycerides | 9.00 | 6.00 | 5.00 | 1.50 | 1.80 |
| Average | 10.50 | 8.50 | 3.75 | 1.29 | 3.33 |

Table 2: **Average $\chi^2$ by association test at Bonferroni significant genes for circulating lipid phenotypes.** The average was taken across the union of genes declared significant by either COAST or SKAT-O. COAST is the coding-variant allelic series test, ALL is SKAT-O applied to all variants, and PTV is SKAT-O applied to protein truncating variants. COAST/ALL is the ratio of COAST to ALL and COAST/PTV is the ratio of COAST to PTV.

| Trait | COAST | ALL | PTV | COAST/ALL | COAST/PTV |
|---|---|---|---|---|---|
| Cholesterol | 124.72 | 97.22 | 63.95 | 1.28 | 1.95 |
| HDL | 119.16 | 106.07 | 29.24 | 1.12 | 4.07 |
| LDL | 184.60 | 150.53 | 102.57 | 1.23 | 1.80 |
| Triglycerides | 111.51 | 83.02 | 78.09 | 1.34 | 1.43 |
| Average | 135.00 | 109.21 | 68.46 | 1.24 | 2.31 |

Table 3: **Overlap with Genebass and the GWAS catalog results of significant circulating lipid genes identified by the coding-variant allelic series test (COAST).** Genebass contains rare-variant associations while the GWAS catalog common-variant associations. $n_{\text{total}}$ is the total number of Bonferroni significant associations. $n_{\text{catalog}}$ is the number of genes associated with the same trait in the GWAS catalog, while $n_{\text{genebass}}$ is the number of genes associated with the same trait in Genebass. $n_{\text{both}}$ is the number of genes associated in both, while $n_{\text{neither}}$ is the number genes associated in neither.

| Trait | $n_{\text{total}}$ | $n_{\text{catalog}}$ | $n_{\text{genebass}}$ | $n_{\text{both}}$ | $n_{\text{neither}}$ |
|---|---|---|---|---|---|
| Cholesterol | 11 | 9 | 11 | 9 | 0 |
| HDL | 15 | 13 | 15 | 13 | 0 |
| LDL | 7 | 6 | 7 | 6 | 0 |
| Triglycerides | 9 | 8 | 7 | 6 | 0 |

Table 4: **Overlap with Genebass and the GWAS catalog of significant circulating lipid genes identified by SKAT-O applied to all variants.** Genebass contains rare-variant associations while the GWAS catalog common-variant associations. $n_{\text{total}}$ is the total number of Bonferroni significant associations. $n_{\text{catalog}}$ is the number of genes associated with the same trait in the GWAS catalog, while $n_{\text{genebass}}$ is the number of genes associated with the same trait in Genebass. $n_{\text{both}}$ is the number of genes associated in both, while $n_{\text{neither}}$ is the number genes associated in neither.

| Trait | $n_{\text{total}}$ | $n_{\text{catalog}}$ | $n_{\text{genebass}}$ | $n_{\text{both}}$ | $n_{\text{neither}}$ |
|---|---|---|---|---|---|
| Cholesterol | 7 | 6 | 7 | 6 | 0 |
| HDL | 14 | 12 | 14 | 12 | 0 |
| LDL | 7 | 6 | 7 | 6 | 0 |
| Triglycerides | 6 | 6 | 5 | 5 | 0 |

#### 2.3 Cell Count Phenotypes

##### 2.3.1 Figures

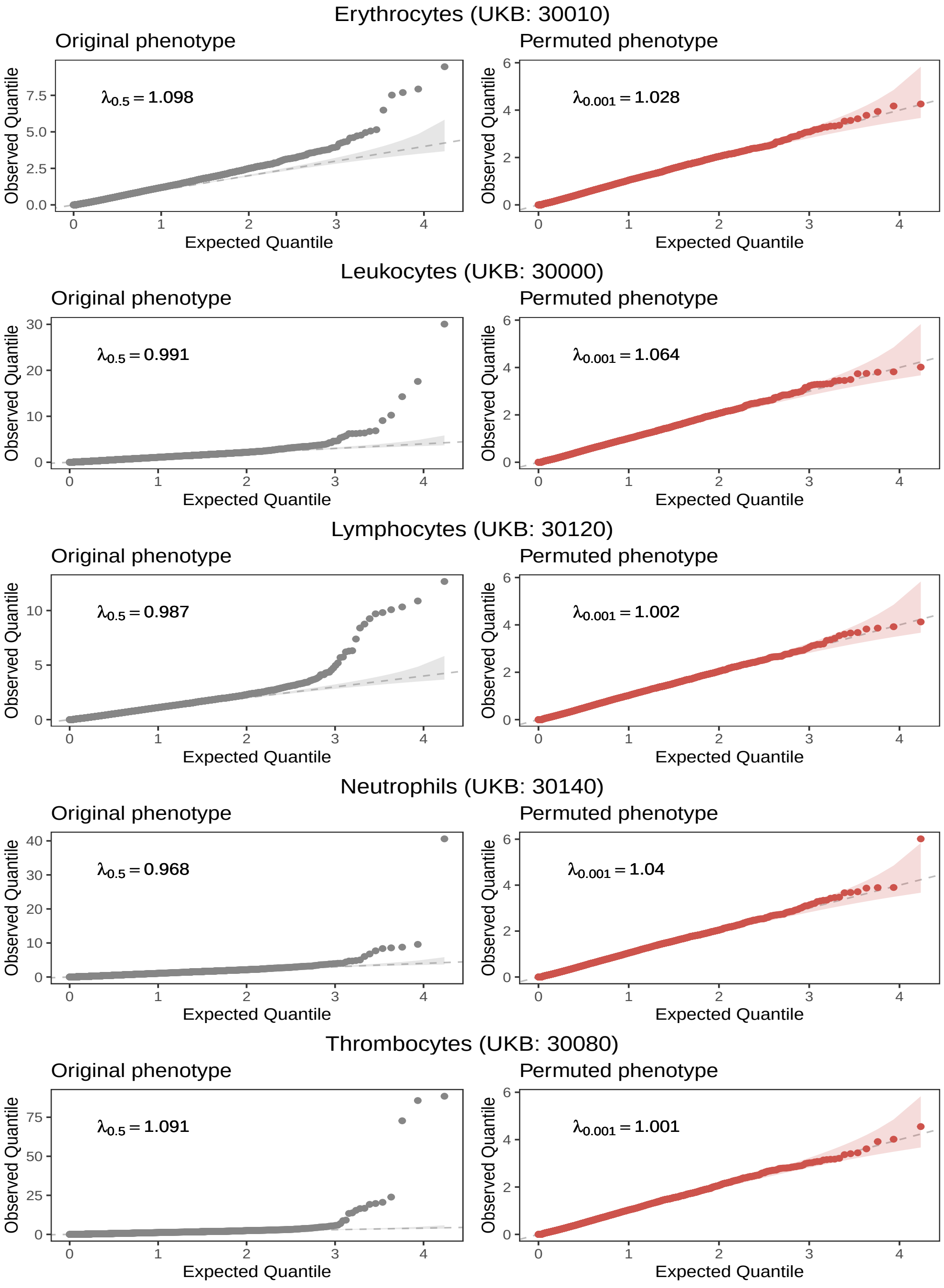

Figure 12: **Uniform quantile-quantile plots for COAST p-values on observed and permuted cell count phenotypes.** $\lambda_p$ is the genomic inflation factor calculated at the $p$th percentile. Note the difference in scale of the Y-axis between the original (left) and permuted phenotype (right).

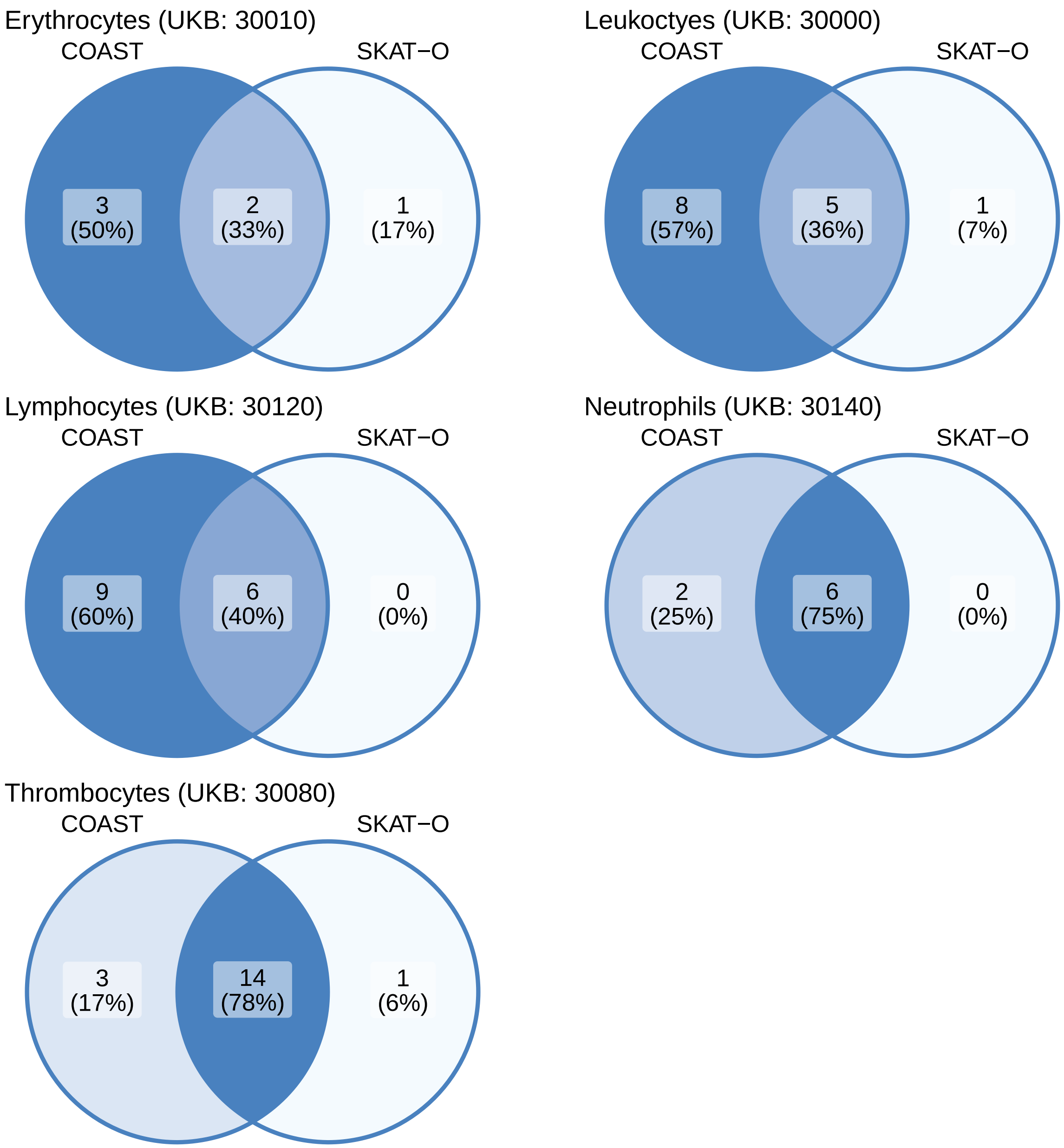

Figure 13: **Venn diagrams for the numbers of Bonferroni significant genes by association test for cell count phenotypes.** COAST is the coding-variant allelic series test. SKAT-O is applied to all rare coding variants (SKAT-O ALL).

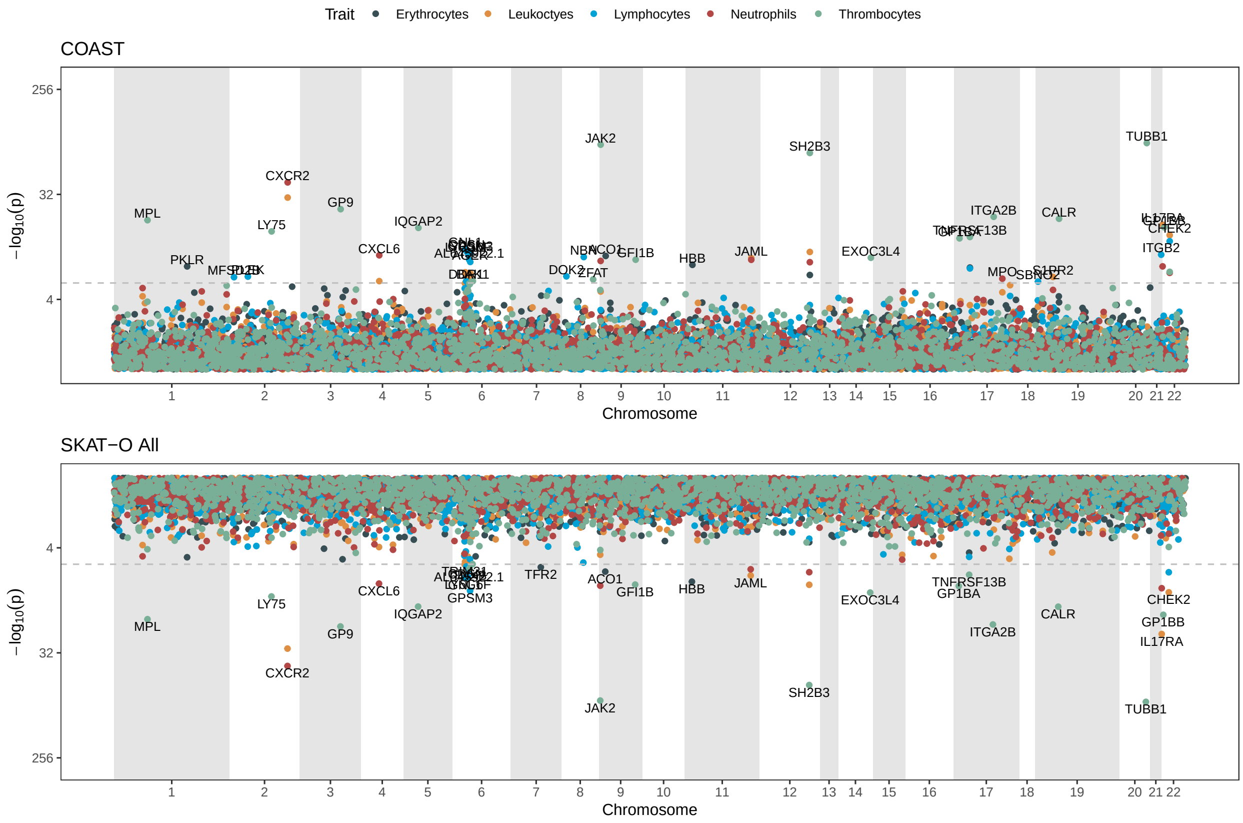

Figure 14: **Mirrored Manhattan plots for cell count phenotypes.** Upper panel is the coding-variant allelic series test (COAST). Lower panel is SKAT-O applied to all rare coding variants (SKAT-O ALL).

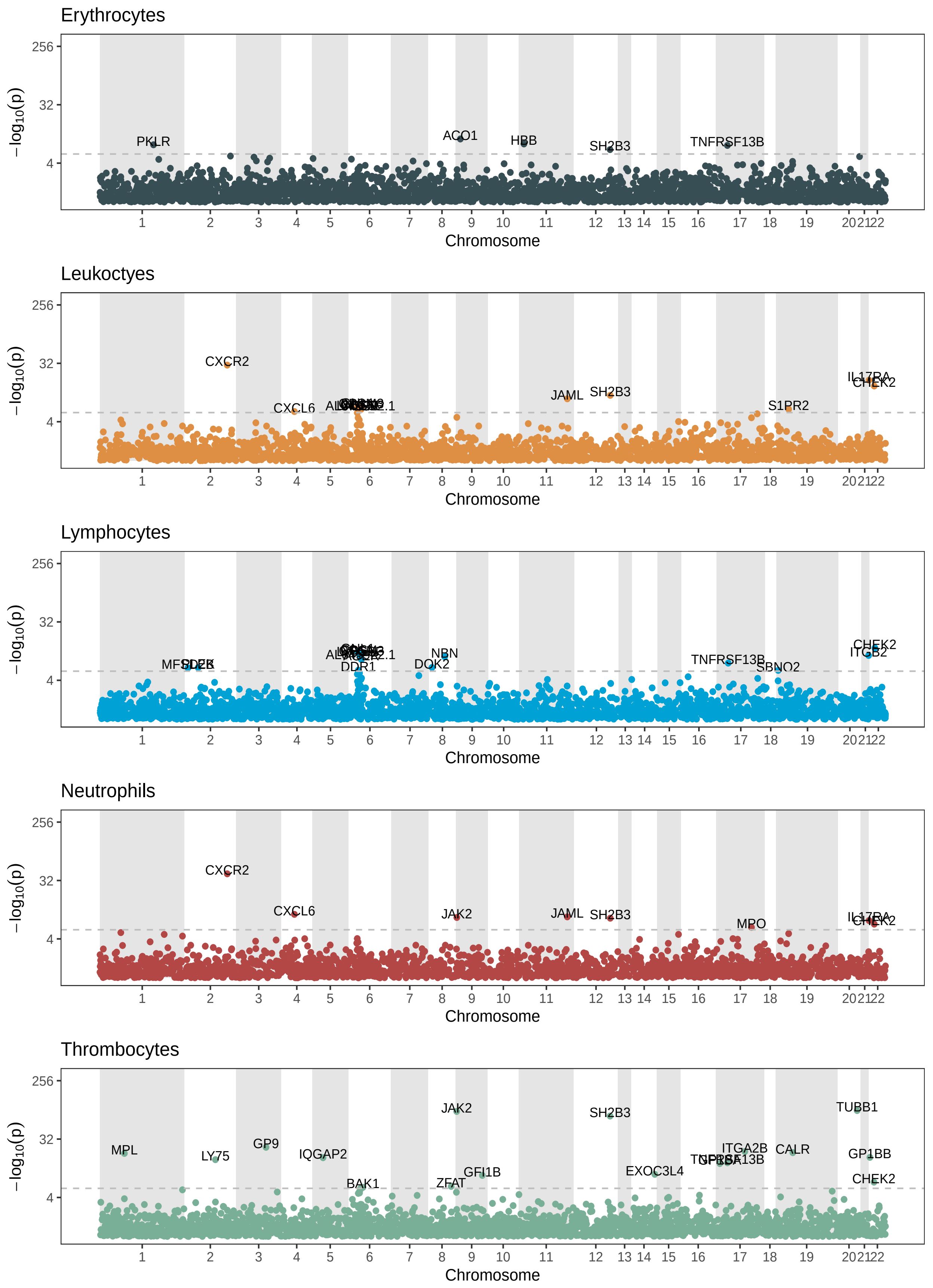

Figure 15: **Stratified Manhattan plots for allelic series (COAST) analysis of cell count phenotypes.**

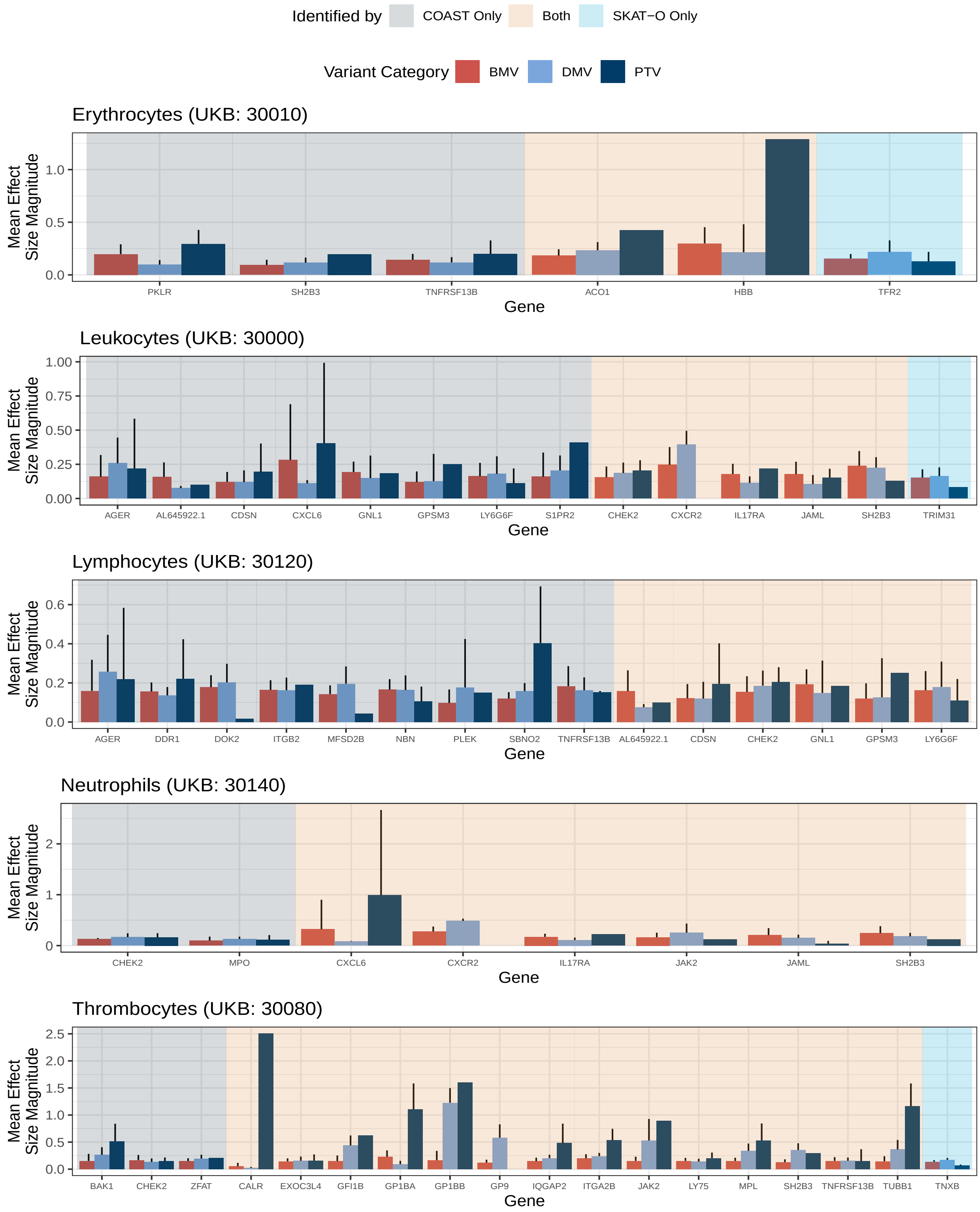

Figure 16: **Effect size patterns among genes significantly associated with cell count phenotypes by the coding-variant allelic series test (COAST).** Effect sizes were estimated by standard linear regression. Each bar represents the mean effect size for variants within a gene and variant category. Error bars are 95% confidence intervals. The absence of an error bar indicates insufficiently many variants to estimate a standard error.

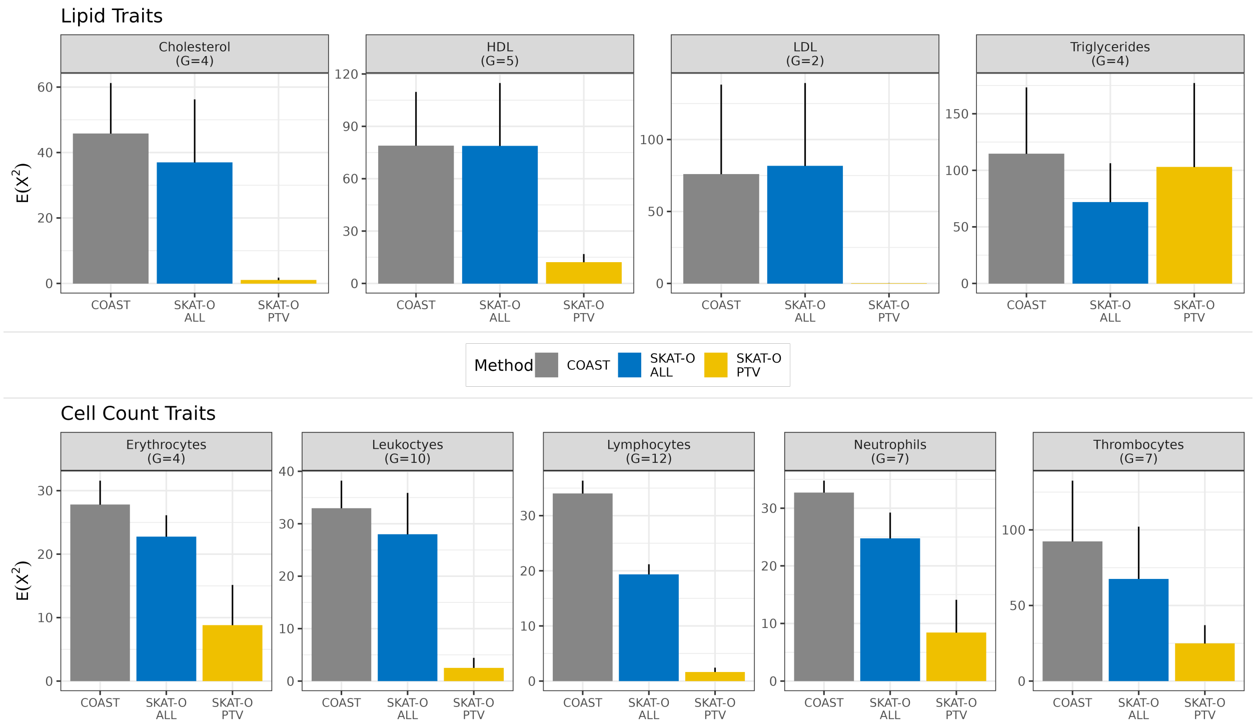

Figure 17: **Average $\chi^2$ statistic at empirical non-allelic series.** A gene was considered a non-allelic series if it was significantly associated with the trait by any association test and the mean effect size for BMVs was greater than the that for DMVs or the mean effect size for DMVs was greater than that for PTVs. Note that the mean effect size is subject to estimation error. The number of non-allelic series represented in each panel is denoted by $G$.

##### 2.3.2 Tables

Table 5: **Counts of Bonferroni significant genes by association test for cell count phenotypes.** COAST is the coding-variant allelic series test, ALL is SKAT-O applied to all variants, and PTV is SKAT-O applied to protein truncating variants. COAST/ALL is the ratio of COAST to ALL and COAST/PTV is the ratio of COAST to PTV.

| Trait | COAST | ALL | PTV | COAST/ALL | COAST/PTV |
|---|---:|---:|---:|---:|---:|
| Erythrocytes | 5.00 | 3.00 | 2.00 | 1.67 | 2.50 |
| Leukoctyes | 12.00 | 5.00 | 1.00 | 2.40 | 12.00 |
| Lymphocytes | 15.00 | 6.00 | 2.00 | 2.50 | 7.50 |
| Neutrophils | 7.00 | 5.00 | 1.00 | 1.40 | 7.00 |
| Thrombocytes | 16.00 | 14.00 | 6.00 | 1.14 | 2.67 |
| Average | 11.00 | 6.60 | 2.40 | 1.82 | 6.33 |

Table 6: **Average $\chi^2$ by association test at Bonferroni significant genes for cell count phenotypes.** The average was taken across the union of genes declared significant by either COAST or SKAT-O. COAST is the coding-variant allelic series test, ALL is SKAT-O applied to all variants, and PTV is SKAT-O applied to protein truncating variants. COAST/ALL is the ratio of COAST to ALL and COAST/PTV is the ratio of COAST to PTV.

| Trait | COAST | ALL | PTV | COAST/ALL | COAST/PTV |
|---|---:|---:|---:|---:|---:|
| Erythrocytes | 27.17 | 19.12 | 11.09 | 1.42 | 2.45 |
| Leukoctyes | 34.06 | 27.55 | 5.24 | 1.24 | 6.50 |
| Lymphocytes | 33.78 | 19.61 | 5.37 | 1.72 | 6.29 |
| Neutrophils | 32.71 | 24.75 | 8.39 | 1.32 | 3.90 |
| Thrombocytes | 110.13 | 94.29 | 23.45 | 1.17 | 4.70 |
| Average | 47.57 | 37.07 | 10.71 | 1.37 | 4.77 |

Table 7: **Overlap with Genebass and the GWAS catalog results of significant cell count genes identified by the allelic series test.** Genebass contains rare-variant associations while the GWAS catalog common-variant associations. $n_{\text{total}}$ is the total number of Bonferroni significant associations. $n_{\text{catalog}}$ is the number of genes associated with the same trait in the GWAS catalog, while $n_{\text{genebass}}$ is the number of genes associated with the same trait in Genebass. $n_{\text{both}}$ is the number of genes associated in both, while $n_{\text{neither}}$ is the number genes associated in neither.

| Trait | $n_{\text{total}}$ | $n_{\text{catalog}}$ | $n_{\text{genebass}}$ | $n_{\text{both}}$ | $n_{\text{neither}}$ |
|---|---|---|---|---|---|
| Erythrocytes | 5 | 5 | 5 | 5 | 0 |
| Leukocytes | 13 | 6 | 13 | 6 | 0 |
| Lymphocytes | 15 | 7 | 14 | 6 | 0 |
| Neutrophils | 8 | 6 | 8 | 6 | 0 |
| Thrombocytes | 17 | 15 | 17 | 15 | 0 |

Table 8: **Overlap with Genebass and the GWAS catalog of significant cell count genes identified by SKAT-O applied to all variants.** Genebass contains rare-variant associations while the GWAS catalog common-variant associations. $n_{\text{total}}$ is the total number of Bonferroni significant associations. $n_{\text{catalog}}$ is the number of genes associated with the same trait in the GWAS catalog, while $n_{\text{genebass}}$ is the number of genes associated with the same trait in Genebass. $n_{\text{both}}$ is the number of genes associated in both, while $n_{\text{neither}}$ is the number genes associated in neither.

| Trait | $n_{\text{total}}$ | $n_{\text{catalog}}$ | $n_{\text{genebass}}$ | $n_{\text{both}}$ | $n_{\text{neither}}$ |
|---|---|---|---|---|---|
| Erythrocyte | 3 | 3 | 3 | 3 | 0 |
| Leukocyte | 6 | 4 | 5 | 3 | 0 |
| Lymphocyte | 6 | 0 | 6 | 0 | 0 |
| Neutrophil | 6 | 5 | 6 | 5 | 0 |
| Thrombocytes | 15 | 14 | 14 | 14 | 1 |
